# SLAMseq resolves the kinetics of maternal and zygotic gene expression in early zebrafish embryogenesis

**DOI:** 10.1101/2022.06.01.494399

**Authors:** Pooja Bhat, Luis E. Cabrera-Quio, Veronika A. Herzog, Nina Fasching, Andrea Pauli, Stefan L. Ameres

## Abstract

The maternal-to-zygotic transition (MZT) is a key developmental process in metazoan embryos that involves the activation of zygotic transcription (ZGA) and degradation of maternal transcripts. We employed metabolic mRNA sequencing (SLAMseq) to deconvolute the compound embryonic transcriptome in zebrafish. While mitochondrial zygotic transcripts prevailed prior to MZT, we uncover the spurious transcription of hundreds of short and intron-poor nuclear genes as early as the 2-cell stage. Upon ZGA, most zygotic transcripts originate from thousands of maternal-zygotic (MZ) genes that are transcribed at rates comparable to those of hundreds of purely zygotic genes and replenish maternal mRNAs at distinct timescales. Rapid replacement of MZ transcripts involves transcript decay features unrelated to major maternal degradation pathways and promotes *de novo* synthesis of the core gene expression machinery by increasing poly(A)-tail length and translation efficiency. SLAMseq hence provides unprecedented insights into the timescales, molecular features and regulation of MZT during zebrafish embryogenesis.

## Introduction

The transition of a fertilized egg to a developing embryo is driven by both maternally inherited (maternal) and zygotically produced (zygotic) factors. After fertilization, the zygotic genome is transcriptionally inactive, and maternally deposited RNA and proteins are essential to carry out all cellular functions (Lee et al., 2014). As development proceeds, control of cellular processes is handed over to transcripts and proteins produced by the zygotic genome. This handover is known as the maternal-to-zygotic transition (MZT), which can last from a few hours in organisms such as *Drosophila*, *C. elegans*, *Xenopus* and zebrafish to days in mouse and humans (Lee et al., 2014; Tadros and Lipshitz, 2009; Vastenhouw et al., 2019). Thus, progression through embryogenesis requires the complete rearrangement of the transcriptome: maternal transcripts need to be cleared in a timely manner, while the onset of nuclear transcription produces new mRNA during zygotic genome activation (ZGA).

In zebrafish embryos, the major wave of ZGA occurs after the first ten rapid cleavage cycles approximately three hours post fertilization. In contrast to the nucleus, the mitochondrial genome is transcribed instantly after fertilization (Heyn et al., 2014). Moreover, low-level of stochastic transcription due to local opening of the chromatin, as well as selective transcription from specific loci, e.g., the multi-copy miR-430 locus, have been reported prior to ZGA in zebrafish (Chan et al., 2019; Lee et al., 2013; Miao et al., 2022; Pálfy et al., 2020; Sato et al., 2019; Stapel et al., 2017; White et al., 2017).

Research investigating the timing and dynamics of ZGA has largely focused on the relatively small fraction of purely zygotic transcripts (estimated to be <25%; (Vastenhouw et al., 2019)), which lack maternal mRNA counterparts and can therefore be directly quantified from steady-state RNA-sequencing experiments (Aanes et al., 2011; Pauli et al., 2012). However, most transcripts encoding housekeeping proteins, including components required for essential cellular processes like transcription, translation, metabolism and homeostasis, are already produced during oogenesis, maternally inherited and thus initially available as templates for *de novo* synthesis of protein products in the embryo before being replaced by newly synthesized mRNAs in the embryo. During a transition period, both maternal (old) and zygotic (new) transcripts are present in the embryo for this group of so-called maternal-zygotic (MZ) genes. MZ genes comprise a potentially large fraction that essentially remains uncharacterized in their transcriptional and post-transcriptional regulation during early embryogenesis due to technical challenges in distinguishing maternal from zygotic contributions particularly for this particular class of genes.

The gap in knowledge with respect to the relative amount and dynamics of MZ gene expression is largely due to difficulties in quantitatively assessing transcripts in a genome-wide and time-of-synthesis-aware manner. While some genes are known to differ in maternal versus zygotic promoter and/or 3′ UTR usage (Bhat et al., 2021; Haberle et al., 2014; Lawson et al., 2020; Ulitsky et al., 2012), this is not a universal feature for all MZ genes. Previous efforts to characterize *de novo* synthesized transcripts in a global manner have employed sequencing-based technologies (Lee et al., 2014), including quantification of intron-containing pre-mRNA (Lee et al., 2013; Rabani et al., 2014), perturbations to inhibit zygotic transcription (Lee et al., 2013), the usage of divergent parental zebrafish strains enabling SNP-mapping of mRNAs transcribed from the paternal genome (Harvey et al., 2013), and metabolic labeling of newly synthesized RNA in the embryo followed by biochemical enrichment of labeled RNA (Heyn et al., 2014). However, none of these methods were able to provide a sufficiently sensitive and comprehensive analysis of the unperturbed embryonic transcriptome in a maternal/zygotic-aware manner. Inhibition of *de novo* transcription causes cellular stress and impacts RNA stability and localization (Bensaude, 2014; Lee et al., 2013; Lugowski et al., 2018); pre-mRNA quantification and SNP-mapping is limited in scope and fails to address the complete zygotic transcriptome; and metabolic labeling coupled to biochemical fractionation is laborious, requires high amounts of starting material and relies on normalization procedures that frequently fail to account for unlabeled transcript contaminants (Lugowski et al., 2018; Schwalb et al., 2016). Given that maternal transcripts are in large excess during the initial stages of embryogenesis, un-labelled maternal contaminants are of particular concern for quantitatively assessing *de novo* synthesized transcripts during ZGA. It has therefore remained unclear how big the fraction of MZ transcripts is relative to the total amount of *de novo* synthesized transcripts. Moreover, the timing of transcription of MZ genes and dynamics of replacement of their maternal with their zygotic transcripts have not been addressed in a quantitative and comprehensive manner.

Beyond the intricacies in the regulation of the zygotic transcriptome, the formation of gene products in embryos is also controlled at the level of translation. Prior to MZT, the efficiency of translation is largely determined by poly(A)-tail length in zebrafish, while later in development (at gastrulation as well as in most non-embryonic cells), control of gene expression is dominated by mRNA production and decay (Subtelny et al., 2014). During early embryogenesis, polyadenylation of mRNAs therefore plays a more direct role in translational regulation, beyond the established effect on mRNA stabilization (Richter, 1999). While poly(A)-tail length is usually determined by nuclear poly(A) polymerases after mRNA 3′ end formation during transcriptional termination, poly(A)-tails can be re-extended in the cytoplasm of early embryos to selectively enhance protein output encoded by maternal mRNA (Aanes et al., 2011; Subtelny et al., 2014; Weill et al., 2012; Winata et al., 2018). Hence, early embryonic development is governed by compound switches in both transcriptional and translational control, but their interplay remains largely elusive.

The rate of RNA replacement during MZT depends both on the rate of *de novo* synthesis of zygotic transcripts and on the rate of maternal mRNA decay. The latter is determined by *cis* elements within transcripts, including codon optimality and 3′UTR length, as well as *trans* regulators, including RNA binding proteins (RBPs) and microRNAs (miRNAs) (Bazzini et al., 2016; Filipowicz et al., 2008; Hentze et al., 2018; Mishima and Tomari, 2016). Two of the best characterized post-transcriptional regulators during early development promote CCR4-NOT-dependent deadenylation of mRNA: The zygotically expressed zebrafish miRNA miR-430 induces degradation of target site containing RNAs, while ribosome slowdown causes co-translational decay of maternal RNA enriched in rare codons (codon-mediated decay) (Bazzini et al., 2016; Giraldez, 2006; Mishima and Tomari, 2016; Mishima et al., 2022). Upon deadenylation, Tut-4/-7-mediated deposition of uridines at the end of short poly(A)-tail-containing mRNAs promotes decay (Chang et al., 2018). While some of the mechanisms described may target both maternal and zygotic transcripts, the dynamics and extent of zygotic RNA decay remain largely elusive.

Here, we employ the recently developed SLAMseq technology to classify and characterize the embryonic zebrafish transcriptome at unprecedented resolution and in a maternal/zygotic-aware manner (Herzog et al., 2017a). SLAMseq overcomes the challenges of prior experimental strategies by (1) labelling *de novo* synthesized RNAs with the uridine analog 4-thiouridine (4sU), and (2) quantitatively converting incorporated 4sU moieties *in vitro* during the reverse transcription step into T>C mismatches, which are readily detected and quantified by standard next generation sequencing (NGS) coupled to nucleotide conversion-aware bioinformatic tools (Herzog et al., 2017a; Neumann et al., 2019). Combined with 3′ mRNA sequencing we refined and expanded current mRNA 3′ annotations during zebrafish embryogenesis and characterized the timing, dynamics and extent of transcriptional output during MZT. Our study provides evidence for the spurious transcription of hundreds of genes before the major wave of ZGA and reveals that the bulk of transcriptional output during early embryogenesis is derived from MZ genes and has so far been largely ignored. Finally, we uncover distinct kinetics of transcript replacement for specific sub-classes of MZ genes with stable steady-state expression patterns, of which transcripts encoding factors of the core gene expression program show particularly rapid exchanges between maternal and zygotic transcripts.

## Results

### *In vivo* SLAMseq distinguishes zygotic from maternal transcripts in zebrafish embryos

To investigate gene expression dynamics during MZT, we established an *in vivo* metabolic RNA sequencing approach in zebrafish (Fig. 1a): We injected zebrafish embryos at the one-cell stage with 4-thiouridine (4sU) under conditions that did not interfere with embryonic development (1.5 mM final concentration of 4sU; Fig. S1a and b) or normal steady-state gene expression (Fig. S1c). Following total RNA extraction at different developmental time points up to 5.5 hours post fertilization (hpf), we performed thiol-linked alkylation to uncover newly synthesized transcripts in cDNA libraries as reported by specific T-to-C (T>C) conversions at the sites of 4sU incorporation (SLAMseq) (Herzog et al., 2017a). We coupled SLAMseq to 3′ mRNA sequencing, which we previously showed to provide rapid and scalable experimental access to the cellular kinetics of RNA polymerase II-dependent gene expression in a transcript 3′ isoform-sensitive manner (Herzog et al., 2017a; Muhar et al., 2018). To overcome constraints imposed by the poor assignment of mRNA 3′ ends in the available zebrafish genome release (GRCz11), we re-annotated stage-specific poly-adenylation sites using 3′GAmES (Fig. S1d and e) (Bhat et al., 2021). When inspected manually, 3′GAmES-derived counting windows overlapped the 3′ end of gene units as delineated by conventional RNAseq (gene body) and CAGE-seq (5′ end) (Fig. 1b). Moreover, transcriptome-wide comparisons of conventional RNAseq with 3′ mRNA end sequencing revealed more even mRNA expression recalls for 3′ GAmES-derived versus Ensembl annotations (Fig. S1f and g). Notably, in contrast to conventional RNAseq of poly(A)-enriched samples, 3′ mRNA sequencing employs short anchored oligo(dT) priming for cDNA library preparation, which diminishes biases in quantification of mRNA with different poly(A)-tail length (Fig. S1h - l). This is particularly important during zebrafish embryogenesis, since poly(A)-tail length-variants frequently emerge from cytoplasmic mRNA de- and re-adenylation (Aanes et al., 2011; Winata et al., 2018).

**Figure 1.**
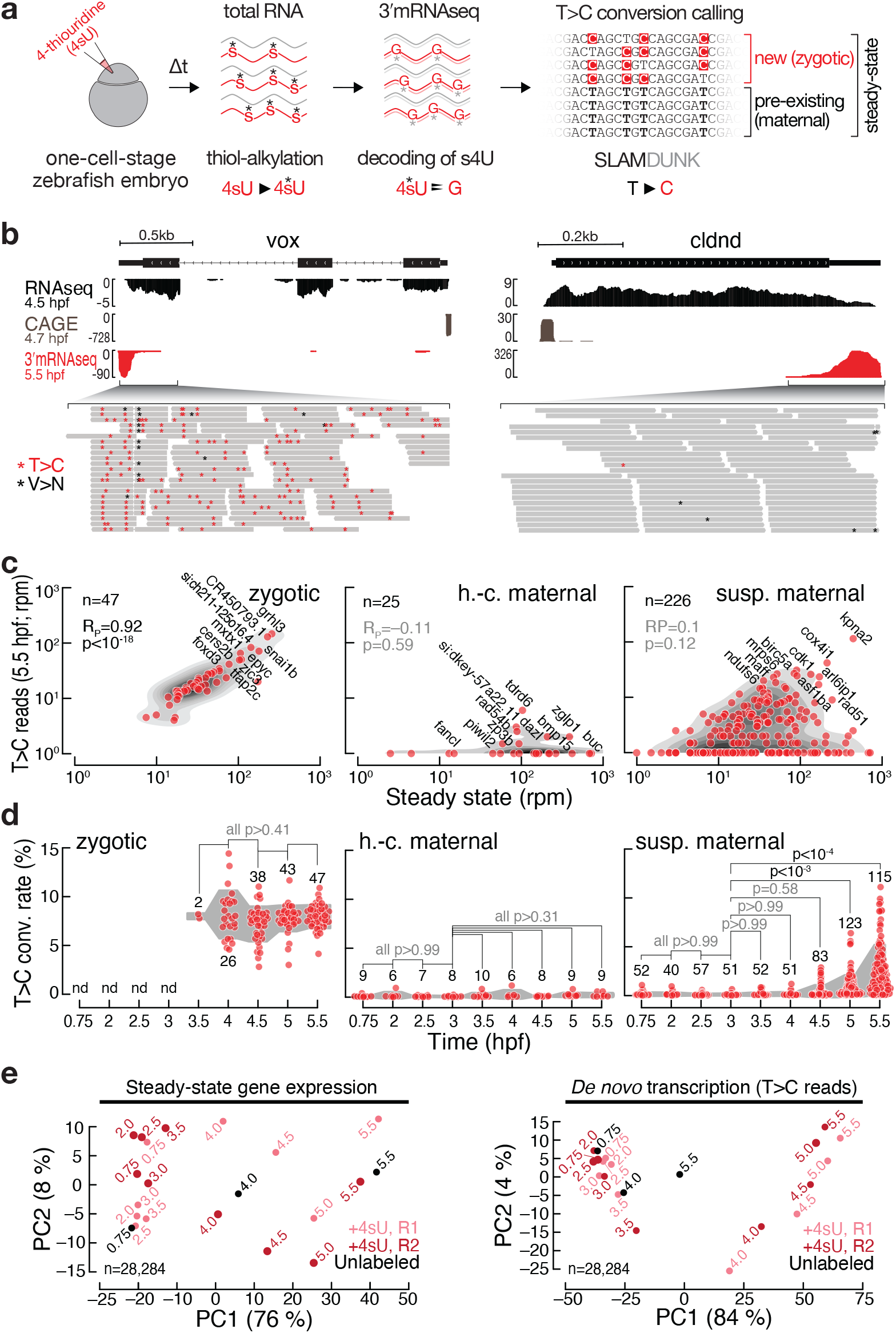
Time-resolved 3ʹ mRNA sequencing (3′mRNA SLAMseq) in zebrafish embryos distinguishes zygotic from maternally deposited mRNA. **(a)** Experimental setup for time-resolved 3′ mRNA sequencing (3′ mRNA SLAMseq) during zebrafish embryogenesis. Zebrafish embryos were injected with 4-thiouridine (4sU; 1.5 mM final conc.) at the one-cell-stage and development was allowed to proceed for a defined time (Δt), followed by total RNA preparation and SLAMseq employing 3′ mRNA sequencing (3′ mRNAseq). *De novo* (i.e., zygotic) transcripts were distinguished from maternal mRNAs by the presence of T>C conversions at sites of 4sU incorporation and a nucleotide conversion-aware data analysis pipeline (SLAMDUNK) (Neumann et al., 2019). **(b)** Representative genome browser screenshot of the known example zygotic gene *ventral homeobox* (*vox*; left) and the maternal gene *claudin d* (*cldnd*; right). Tracks for poly(A)-selected RNAseq (top), cap analysis gene expression (CAGE; middle; remapped from (Haberle et al., 2014)) and 3′ mRNA SLAMseq (bottom), sampled at the indicated developmental stage (hours post fertilization; hpf), are shown. Individual reads of 3′mRNA SLAMseq datasets mapping to the respective 3′ UTR counting windows are shown in the zoom-in. Asterisks indicate T>C conversions (T>C; red) or any other conversions (V>N; black). **(c)** Comparison of *de novo* transcription (T>C reads in reads per million; rpm) and steady-state gene expression (steady-state in rpm) for pre-defined zygotic (n=47), high-confidence (h.-c.) maternal (n=25) and suspected (susp.) maternal genes (n=226) at 5.5 hpf. Pearson correlation coefficient (R_P_) and associated p-values (p) are shown. Abbreviations of example genes are indicated. **(d)** T>C conversion (conv.) rate for transcripts defined in (c) during MZT (time in hours post fertilization, hpf). Individual genes for which a confident average conversion rate could be derived from two independent biological replicates (red, n = number of genes) and interquartile range (grey area) are shown. P-values (p; Kruskal-Wallis and Dunn’s multiple comparison test) are indicated; nd, not detected. **(e)** Principal component analyses of unfiltered control (unlabeled; black) and two independent 3′ mRNA SLAMseq (+4sU; R1 light and R2 dark red) experiments at the indicated developmental timepoints (in hours post fertilization). Steady-state gene expression (left); *de novo* transcription (T>C conversion containing reads) (right). The number of genes (n) considered for the analysis is shown. The variance of each principal component is indicated in percent.

To test if time-resolved 3′ mRNA SLAMseq distinguishes newly synthesized zygotic from maternal transcripts, we inspected libraries generated from zebrafish embryos 5.5 hpf – a timepoint at which robust ZGA had already occurred. As expected, we observed a strong accumulation of T>C conversions in reads mapping to the known zygotic gene *vox* but not the maternal transcript *cldnd* (Fig. 1b). To assess the specificity and sensitivity of SLAMseq, we inspected a broader set of pre-defined zygotic (i.e. detected by Harvey *et al*., 2013, Heyn *et al*., 2013 or Lee *et al*., 2013, and further filtered; see Materials and Methods; n=47), high-confidence (h.-c.) maternal (i.e. genes involved in oogenesis and meiosis and not expressed in somatic adult tissues; n=25) and suspected (susp.) maternal transcripts (i.e. previously defined by Lee et al., 2013 and further filtered, see Materials and Methods; n=226) (see Materials and Methods and Fig. S1m): At 5.5 hpf, *de novo* transcription correlated significantly with steady-state expression for zygotic genes (Pearson correlation R_P_=0.92, p<10^-18^, with ∼52% of all reads exhibiting T>C conversions) but not for h.-c. maternal (R_P_=-0.11, p=0.59) or susp. maternal genes (R_P_=0.1, p=0.12) (Fig. 1c). Notably, we observed a dual distribution in the case of susp. maternal genes, where many genes (n=133) showed signals of *de novo* expression, while others (n=93) lacked a labeling signal, resembling h.-c. maternal genes (Fig. 1c, right plot, S1n, o). To assess the contribution of zygotic transcription to overall steady-state transcript abundances across MZT, we computed T>C conversion rates per gene after accounting for genotype-specific SNPs and sequencing error (see Materials and Methods for details). We found that for zygotic genes – once robustly expressed (i.e., detected in two independent replicates) – the T>C conversion rate per gene reached an immediate median of ∼7.9% (i.e., one in every 13 uridines is replaced by 4sU) and the T>C conversion rate stayed constant for the duration of the experiment (i.e., up to 5.5 hpf), preempting concerns about possible 4sU dilution effects (Fig. 1d, left plot). Note, that T>C conversions did not accumulate in uninjected samples, confirming the overall high signal-to-noise ratio in the assay (Fig. S1p). High-confidence maternal transcripts generally exhibited very low to undetectable T>C conversion rates that did not increase significantly throughout the time-course, revealing a lack of transcription at these genomic loci (Fig. 1d, central plot). In contrast, suspected maternal transcripts displayed constantly low T>C conversion rates at early timepoints, but a significant and continuous increase after 4.5 hpf (Fig. 1d, right plot), which indicates a gradual replacement of select maternal transcripts by their zygotic counterparts despite an overall decrease in steady-state expression (Fig. S1m). Our data therefore shows that it is insufficient to classify maternal transcripts merely by their decrease in steady-state expression, and that a substantial fraction of previously categorized maternal genes may in fact be re-expressed during MZT.

To assess whether 3′ mRNA SLAMseq can reproducibly distinguish stages before and after MZT, we performed principal component analysis of steady-state gene expression across the entire time-series. This analysis showed that the data clustered well for 4sU-injected replicates and the respective control samples, where 76% of variation could be explained by the first principal component which reports on developmental timing (Fig. 1e, left plot). Similar analyses on unfiltered T>C reads resulted in a stronger separation based on 4sU labelling (84% of variation across the first principal component), where 4sU-injected replicates cluster together with control samples prior, but reproducibly distinct from their respective unlabeled controls post MZT (Fig. 1e, right plot). Together with the fact that replicate data showed a strong and significant correlation for steady-state gene expression across all timepoints (Pearson correlation R>0.92, p<10^-15^; Fig. S1q, left plots) and *de novo* transcription (assessed conservatively by the presence of multiple T>C conversions per read) as early as 2.5 hpf (R_P_>0.89, p<10^-12^; Fig. S1q, right plots), we conclude that SLAMseq can reproducibly distinguish stages before and after MZT. Notably, we obtained similar results in experiments where embryos were incubated with 4sU rather than subjected to injection, albeit with ∼10-fold lower labeling efficiencies (data not shown).

### The early embryonic transcriptome is dominated by pre-existing maternal mRNA

Though several studies have attempted to systematically characterize the timing and extent of transcription during MZT in zebrafish, the overall scope and contribution of zygotic gene expression to the steady-state transcriptome remains largely unknown. To assess transcriptional activity before and after the major wave of ZGA, we determined the number of T>C reads in 3′ mRNA SLAMseq datasets (T>C reads; +4sU) relative to unlabeled samples (–4sU; Fig. 2a) or in-sample background conversions (i.e., T>A; Fig. S2a). Already at the earliest timepoint (i.e., 0.75 hpf) and much more prominently later (i.e., 5.5 hpf), we detected a significantly higher number of T>C conversion containing reads in 3′ mRNA SLAMseq datasets prepared from zebrafish embryos injected with 4sU (+4sU) than in unlabeled samples (–4sU) or in-sample control conversions (T>A, Whitney Mann p<10^-15^) (Fig. 2a and S2a). To determine the transcriptional output along the entire time course, we determined the number of reads with 1, 2 or ≥3 T>C conversion(s) mapping to the mitochondrial or the nuclear genome (Fig. 2b). Note, that multiple T>C conversions are a confident indicator of *de novo* transcription, as confirmed by the assessment of zygotic and high-confidence maternal transcripts (Fig. S2b). While we detected a continuous high transcription signal in mitochondrial transcripts as early as 0.75 hpf, genes encoded by the nuclear genome exhibited little evidence for robust transcription until 3.5 hpf, followed by a strong increase in all following timepoints (Fig. 2b). We concluded that in contrast to the nuclear genome, robust transcription in mitochondria occurs instantly after fertilization.

**Figure 2.**
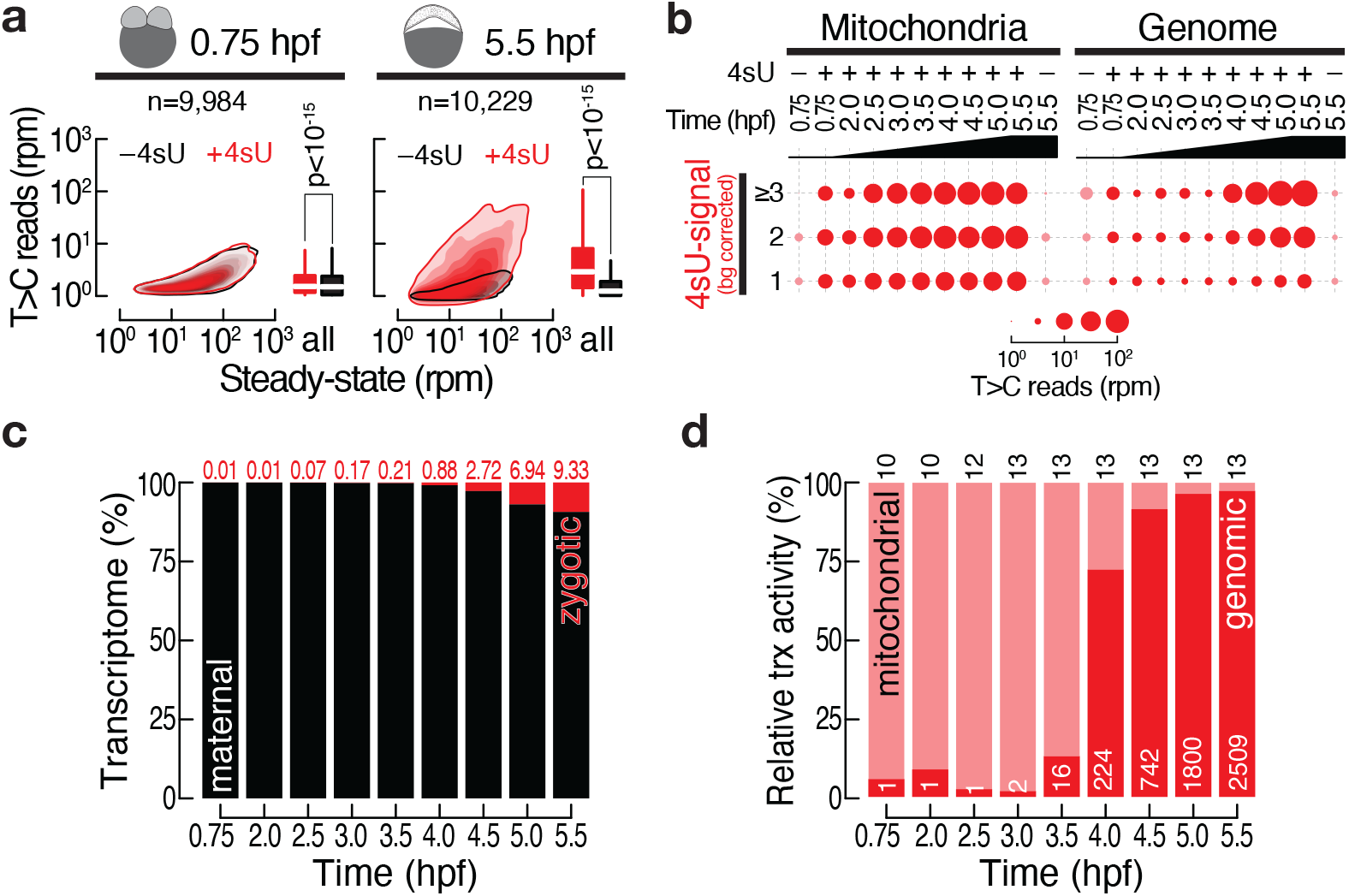
The transcriptional landscape in zebrafish embryogenesis. **(a)** Contour plots show a comparison of *de novo* transcription (i.e., T>C conversion containing reads in rpm) with steady-state gene expression (in rpm) in zebrafish embryos injected with 4sU (+4sU, red) or in untreated control embryos (−4sU, black) as determined by 3′ mRNA SLAMseq at the indicated developmental time points (in hours post fertilization, hpf). Boxplots show a quantitative comparison of *de novo* transcription signal by comparing T>C conversion containing reads. P-value was determined by Wilcoxon test. The number of inspected genes is indicated (n). **(b)** Balloon plots report the number of reads (in rpm) containing one, two, or three and more T>C conversions mapping to the mitochondrial (left) or the nuclear genome (right) in 3′ mRNA SLAMseq datasets generated from zebrafish embryos injected with 4sU (+4sU) or untreated (–4sU) at the indicated timepoints (in hpf). **(c)** The contribution of zygotic (red, percentage indicated on top) and maternal (black) transcripts to steady-state gene expression at the respective time points (hpf) was determined by quantifying the number of T>C conversion containing transcripts in 3′ mRNA SLAMseq datasets normalized to overall labeling-efficiency and relative to steady-state. For any given gene, *de novo* transcripts were accounted for only if the T>C conversion signature was significantly higher (99% confidence cutoff) compared to unlabeled control samples at any given time point and thereafter. **(d)** Relative transcriptional activity (as determined in (c)) of the mitochondrial (light red) and nuclear genome (dark red) at the indicated time points (in hpf). The number of genes for each category is indicated.

To estimate the zygotic contribution to the overall transcriptome throughout embryogenesis, we quantified the transcriptional output by determining the overall number of T>C conversion-containing reads (adjusted for labeling efficiency, see Materials and Methods) relative to steady-state (Fig. 2c). We limited this analysis to robustly expressed genes, whose T>C conversion signature (see Fig. S2a) is significantly higher compared to unlabeled control samples (i.e., above 99% confidence cutoff at any given timepoint and thereafter). We found that only 0.01% of the transcriptome comprised newly synthesized gene products at the earliest time points, which only slightly increased prior to the major wave of ZGA. Notably, even at 5.5 hpf only 9.33% of the transcriptome could be assigned to the zygotic genome (Fig. 2c). This suggests that even at the onset of gastrulation the transcriptome is largely dominated by pre-existing maternal gene products. The low levels of zygotic transcripts observed prior to ZGA largely originated from the mitochondrial genome (≥ 85% of the nascent transcriptome), while the nuclear genome dominated the zygotic transcriptome only after 3.5 hpf (Fig. 2d).

### Evidence for the spurious transcription at hundreds of gene loci prior to the major wave of ZGA

Stochastic gene expression is a hallmark of early ZGA (Chan et al., 2019; Heyn et al., 2014; Kwasnieski et al., 2019; Stapel et al., 2017). In fact, recent imaging studies revealed spurious expression of eight genes (*apoeb*, *aldob*, *slc25a22*, *mex3b*, *lrwd1*, *sox19a*, *zic2b* and *tbx16*) at the 512-cell stage (Stapel et al., 2017), and the multi-copy *miR-430* locus as early as the 64-cell-stage (Chan et al., 2019). Given the specificity of 3′ mRNA SLAMseq, we revisited the timing and scope of spurious nuclear transcription at the 2-cell (0.75 hpf) and the 64-cell stage (2 hpf). To this end, we applied a filtering parameter that is based on the rate of T>C conversions (PTC) and reports on the occurrence of individual 4sU incorporation events within transcripts (see Materials and Methods for details). We defined replicate-specific PTC-cutoffs based on the detection of most pre-defined zygotic but no high-confidence maternal genes (Fig. S3a). By applying this filtering scheme, we identified a total of 501 (0.75 hpf) and 554 (2 hpf) genes that were detected above PTC-cutoff, including eight out of the nine previously described early expressed genes (Fig 3a). The number of genes above PTC-cutoff increased substantially to 3.328 genes at 5.5 hpf (Fig. 3b, c and S3b). Notably, only 4.2% (0.75 hpf) and 7.6% (2 hpf) of genes overlapped between replicates at the 2-cell and 64-cell stage, respectively (Fig. 3b), perhaps reflecting a spurious and/or stochastic pattern of expression (Chan et al., 2019; Stapel et al., 2017). Consistent with this, only 20.2% and 18.9% of genes detected at 0.75 hpf and 2 hpf, respectively, were also detected in at least one replicate of each following time point (Fig. 3c and S3c). In contrast, >40% of genes were reproducibly detected above cutoff at 5 hpf and 5.5 hpf, attesting a robust activation of the genome post-MZT (Fig. 3b).

**Figure 3.**
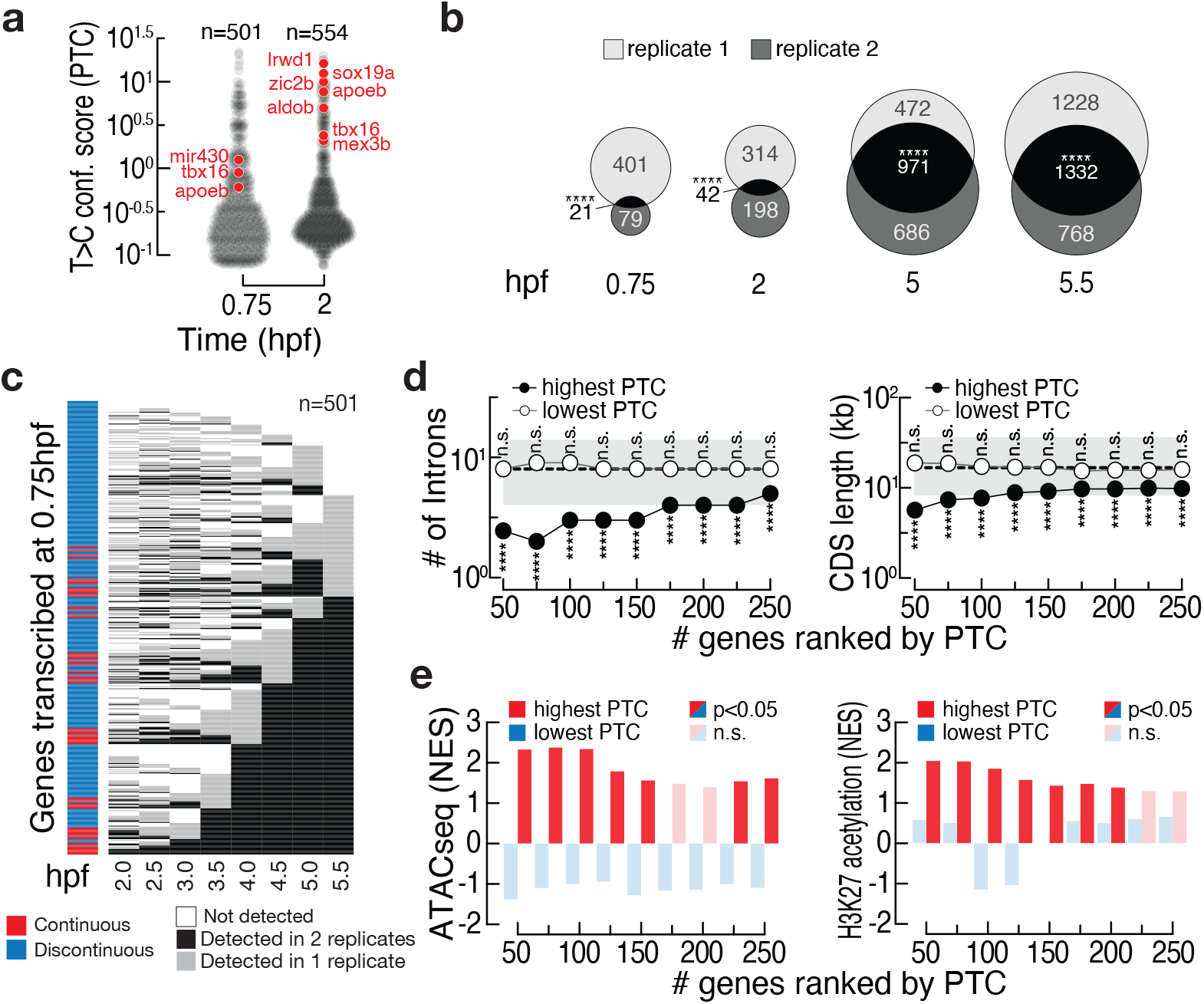
Evidence for the spurious transcription at hundreds of gene loci prior to MZT. **(a)** T>C confidence score (PTC) reports the percent of T>C conversions per 4sU-labeled transcript in percent. Only genes that exhibit a PTC score corresponding to a true positive rate of >75% and false positive rate of 0% (see Fig. S3d) are shown. The number of genes is indicated, and genes previously shown to be transcribed in early zebrafish embryogenesis are highlighted (red). **(b)** Venn diagrams report the overlap of genes detected above PTC-cutoff (see (a)) in two independent biological 3′ mRNA SLAMseq replicates at the indicated time points (in hpf). Significance of overlap was determined by Fisher’s exact test. **** = p<10^-4^. **(c)** Heatmap reports for all genes detected above PTC-cutoff at 0.75 hpf (2-cell stage; n=501) reports the presence of T>C conversion containing reads (above PTC-cutoff) in none (white), one (grey) or both (black) of two independent 3′ mRNA SLAMseq experiments at the indicated subsequent time points (in hpf). Genes with continuous (red) or discontinuous signal (blue) in at least one of two replicates throughout the entire time course are highlighted (left). **(d)** Median number of introns (left) or length of the coding sequence (CDS in kilobases, kb) for the indicated number of genes ranked by highest (filled circles) or lowest (open circles) T>C confidence score (PTC) in 3′ mRNA SLAMseq datasets at 0.75 hpf. Statistically significant deviation (Wilcoxon test; ****=p<10^-4^) from median (dashed line) and interquartile range (grey area) of all genes in the dataset is indicated. **(e)** Normalized enrichment scores (NES) of ATACseq (left; derived from Bogdanović *et al*., 2012) and H3K27 acetylation chromatin immunoprecipitation and sequencing (right; derived from Pálfy *et al*., 2020) for the indicated number of genes ranked by highest (red) or lowest (blue) T>C confidence score (PTC) in 3′ mRNA SLAMseq datasets at 0.75 hpf. Significant enrichment (hypergeometric test with Benjamini-Hochberg correction; p<0.05) is indicated in opaque colors.

In zebrafish and flies, the earliest detected zygotic transcripts are short and contain few introns (Heyn et al., 2014; Kwasnieski et al., 2019). To determine if those features are hallmarks of our earliest-detected gene products, we compared the number of introns and the length of the coding sequence (CDS) of these genes that were grouped based on their PTC ranking at 0.75 hpf or 2 hpf. Indeed, the number of introns and the length of the CDS was significantly lower for genes with high T>C confidence score when compared to all annotated genes (p<10^−4^, Wilcoxon test, Fig. 3d and S3d). We also inspected published chromatin features, such as overall accessibility (measured by ATACseq at the 256-cell stage) and H3K27 acetylation (measured by chromatin-immunoprecipitation and sequencing at the dome stage) at the promoters of genes detected above PTC-cutoff (Bogdanović et al., 2012; Pálfy et al., 2020). Indeed, we found that genes with highest PTC were significantly enriched for open chromatin and H3K27 acetylation (hypergeometric test, p<0.05; Fig 3e and S3e). Taken together, our data revealed hundreds of short genes depleted of introns to be spuriously expressed as early as at the 2-cell stage during zebrafish embryogenesis.

### Classification of the MZT transcriptome reveals the zygotic re-expression of most maternal genes

During MZT, the transcriptome consists of gene products that are strictly maternally provided (M), exclusively expressed by the zygote (Z), or maternally provided and re-expressed in the zygote (MZ). While current approaches can confidently classify transcripts as strictly zygotic, the extent of re-expression of maternal genes remains largely unknown. We therefore classified the transcriptome measured by 3′ mRNA SLAMseq based on (1) steady-state gene expression and (2) the ratio of *de novo* synthesized versus steady-state transcripts on a gene-by-gene basis (Fig. S4a). By considering only genes with robust steady-state expression, we categorized the transcriptome into four groups: (1) M-unstable (n=1,416), with decreasing gene expression during MZT and no detectable T>C conversions; (2) M-stable (n=1,569), with constant gene expression during the entire time course and no detectable T>C conversions; (3) Z (n=434) with increasing gene expression, <1 RPM steady-state expression at 2-cell and 64-cell stage, and T>C conversions detected; and (4) MZ gene (n=6,923), with decreasing, constant or increasing gene expression, ≥1 rpm steady-state expression at 2-cell and 64-cell stage, and T>C conversions detected (Fig. 4a). Notably, the vast majority (i.e., 70%) of M genes were re-expressed zygotically during MZT. To independently validate this classification, we compared steady-state gene expression of each group in fertilized oocytes and 2-cell stage embryos. As expected, we observed a statistically significant high correlation for M-stable, M-unstable and MZ genes in oocytes and 2-cell embryos (Pearson correlation R>0.87; p<10^-15^), while Z genes were not detected in oocytes and were still poorly expressed (if at all) in 2-cell stage embryos (Fig. S4b). Gene ontology (GO) term analysis revealed a significant association of M-unstable genes with reproductive processes, M-stable genes with metabolic and other house-keeping activities and Z genes with functions in development and differentiation (Fig. 4b). Notably, MZ genes were significantly associated with processes related to gene expression – including RNA metabolism, processing and biosynthesis (Fig. 4b).

**Figure 4.**
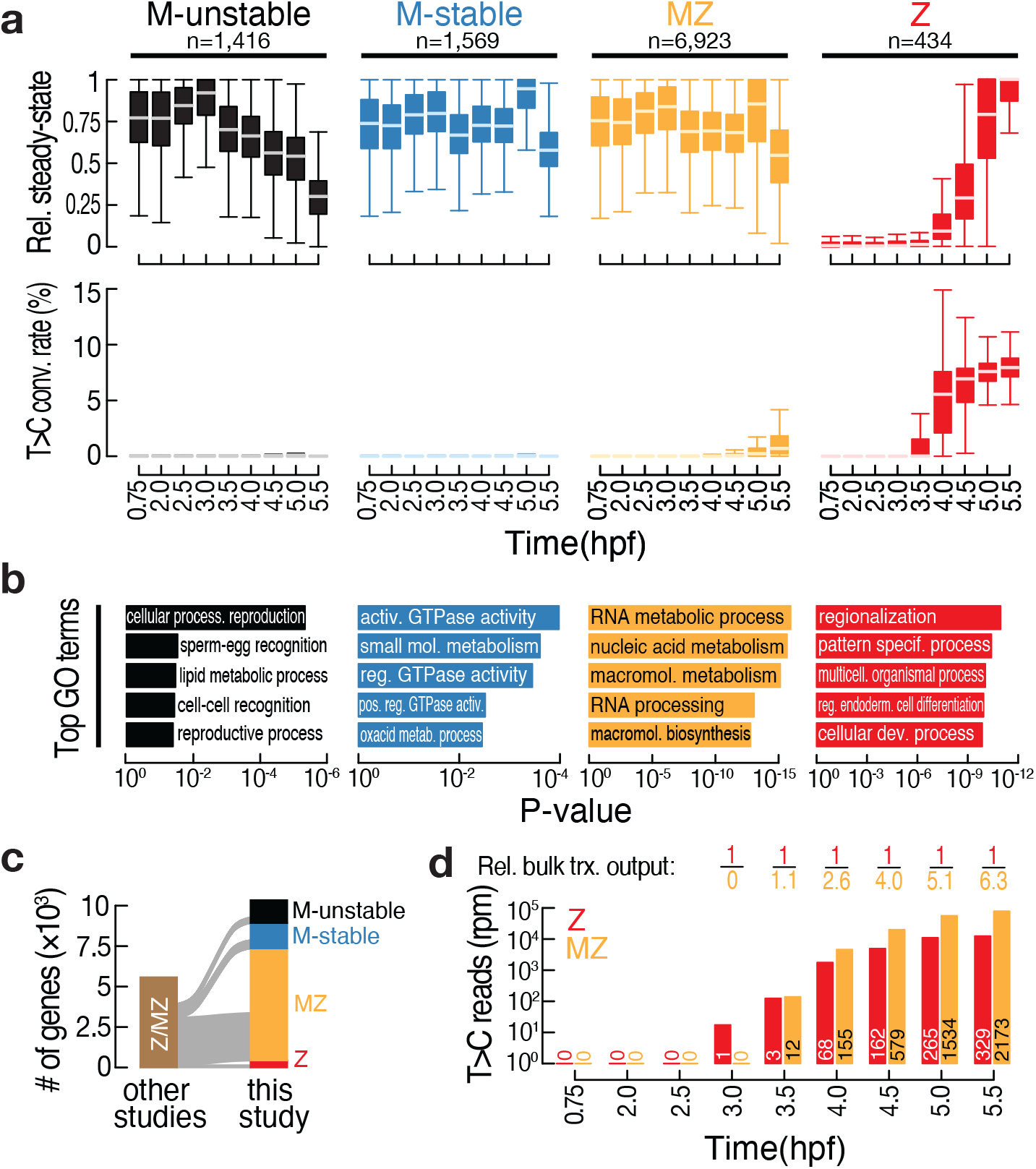
Classification of the zebrafish transcriptome during MZT reveals zygotic re-expression of most maternal genes. **(a)** Classification of the transcriptome by *k*-medioids clustering (see Fig. S4a) was based on steady-state gene expression (top) and T>C conversion rates (bottom), as determined by 3′ mRNA SLAMseq at the indicated developmental time points (in hours post fertilization; hpf). Only genes with a steady-state expression cutoff (> 5 rpm) at any time point were considered. The indicated number of genes (n) were classified into maternal-unstable (M-unstable; black), maternal-stable (M-stable; blue), maternal-zygotic (MZ; yellow) and zygotic (Z; red). Steady-state gene expression was calculated for each gene relative to the maximum observed expression (in rpm). **(b)** Top five gene ontology (GO) terms significantly enriched in each of the gene classes described in (a). The associated p-value is indicated. **(c)** Sankey diagram reports a comparison of 3′ mRNA SLAMseq-based gene classifications (this study) with previously reported zygotic (i.e., zygotic, Z or maternal-zygotic, MZ) gene annotations (Harvey et al., 2013; Heyn et al., 2014; Lee et al., 2013). For further details see Fig. S3c. **(d)** Quantification of bulk zygotic expression (i.e., T>C conversion containing reads in rpm) of zygotic (Z; red) or maternal-zygotic (MZ; yellow) genes, as classified in (a), at the indicated time points (in hpf). The number of genes considered at each time point is indicated. The relative (rel.) bulk transcriptional (trx.) output reports the relative abundance of T>C conversion containing reads derived from all zygotic (red) or maternal-zygotic (yellow) genes.

To benchmark our classification, we compared genes identified to be expressed in zygotes (Z and MZ) by 3′ mRNA SLAMseq to three previous approaches dedicated to the annotation of zygotic transcripts (Harvey et al., 2013; Heyn et al., 2014; Lee et al., 2013) (Fig. S4c): Out of a total of 318 genes reported to be zygotically expressed by metabolic labeling and biochemical enrichment followed by RNA sequencing, we could validate 82.3% (22.3% Z and 60% MZ), while 2.5% were classified as maternal (0.6% M-unstable and 1.8% M-stable) and 15.1% remained below detection limit (i.e. <5RPM) in 3′ mRNA SLAMseq datasets (Heyn et al., 2014). Out of a total of 2,936 genes with assigned zygotic expression by paternal SNP-calling, we could validate 67.8% (66.8% MZ and 1.9% Z), while 15.4% were classified as maternal (5.8% M-unstable and 9.6% M-stable) and 15.9% were undetected (Harvey et al., 2013). And out of a total of 1,706 zygotically expressed genes determined by sequencing of intron-containing pre-mRNA, we could validate 70.13% (59.7% MZ and 10.43% Z), while 1.4% were classified as maternal (0.5% M-unstable and 0.9% M-stable) and 28.4% remained below the chosen cut-off (Lee et al., 2013). Most importantly, beyond the high overall agreement with previous studies, 3′ mRNA SLAMseq revealed 4,398 new zygotically expressed genes (4,188 MZ and 210 Z), which had not been reported previously (Fig. 4c). Among those newly classified genes, both MZ and Z genes exhibited a metabolic sequencing signature that mirrored that observed for pre-defined Z and MZ genes (Fig. S4d, 1d and S1o), further supporting the validity of our classification.

Finally, we compared the extent to which MZ and Z genes contributed to the overall transcriptional load during MZT (Fig. 4d). While robust expression of the first Z gene at 3 hpf – the miR-430 locus – preceded that of the earliest MZ genes (detected at 3.5 hpf), the bulk expression derived from MZ genes outnumbered that of Z genes by up to 6.3-fold. At the same time, the overall higher transcriptional load observed for MZ genes distributed among up to 6.6-times more genes (i.e., 2,173 MZ vs. 329 Z genes at 5.5 hpf; Fig. 4d), resulting in a similar median transcriptional output per locus for MZ and Z genes (Fig. S4e). Taken together, we show that 3′ mRNA SLAMseq deconvolutes the maternal and zygotic origin of transcripts during MZT in zebrafish embryos and uncovers the re-expression of an unexpectedly large number of maternally expressed genes. In fact, we provide evidence that most of the zygotic transcriptome originates from thousands of MZ genes that are on average transcribed at a rate comparable to that of zygotic genes.

### Replacement of maternal gene products during MZT occurs at distinct rates

Given the broad scope of maternal gene activity during MZT (Fig. 4), we asked to what extent newly synthesized transcripts contribute to the steady-state expression of individual genes. To this end, we focused on the subset of MZ genes that did not change in steady-state gene expression during our time-course (i.e., 4,249 out of 6,923 MZ genes) and performed *k*-medioids clustering based on the contribution of T>C conversion-containing reads at multiple time points along the major wave of ZGA (i.e., 3.5 to 5.5 hpf; Fig. 5). As exemplified by the representative genes *dhx15*, *nono*, *pou5f3* and *sox11b*, we could identify four clusters that differed in the rate at which zygotic transcripts had replaced pre-existing maternal gene products (Fig. 5a). Quantitative assessment revealed that cluster 1 (2,268 genes) exhibited only a minor average zygotic contribution of up to 5.29%. The degrees of replenishment were larger for cluster 2 (1,022 genes; up to 15.1%) and cluster 3 (642 genes; up to 30.53%). Genes in cluster 4 (317 genes) were replaced most effectively, with an average zygotic contribution of up to 59.8%, indicating that the steady-state expression of those genes is rapidly dominated by zygotic gene products (Fig. 5b and S5a). Note, that the steady-state expression was significantly lower in cluster 3 and 4 genes as opposed to cluster 1 and 2, albeit all groups exhibited considerably high levels of expression (i.e., median >10 rpm) (Fig. S5b). Since increasing replacement kinetics imply a decrease in transcript stability, we tested if low transcript stabilities are an inherently conserved feature of those (and in particular of cluster 4) genes. To address this, we compared mRNA half-lives of gene cluster orthologs in mouse embryonic stem cells (mESC), as previously determined (Herzog et al., 2017a). In fact, we observed a gradual trend towards decreasing median mRNA half-lives that were significantly lower in genes belonging to cluster 4 (Wilcoxon test, p<10^-2^; Fig. S5c).

**Figure 5.**
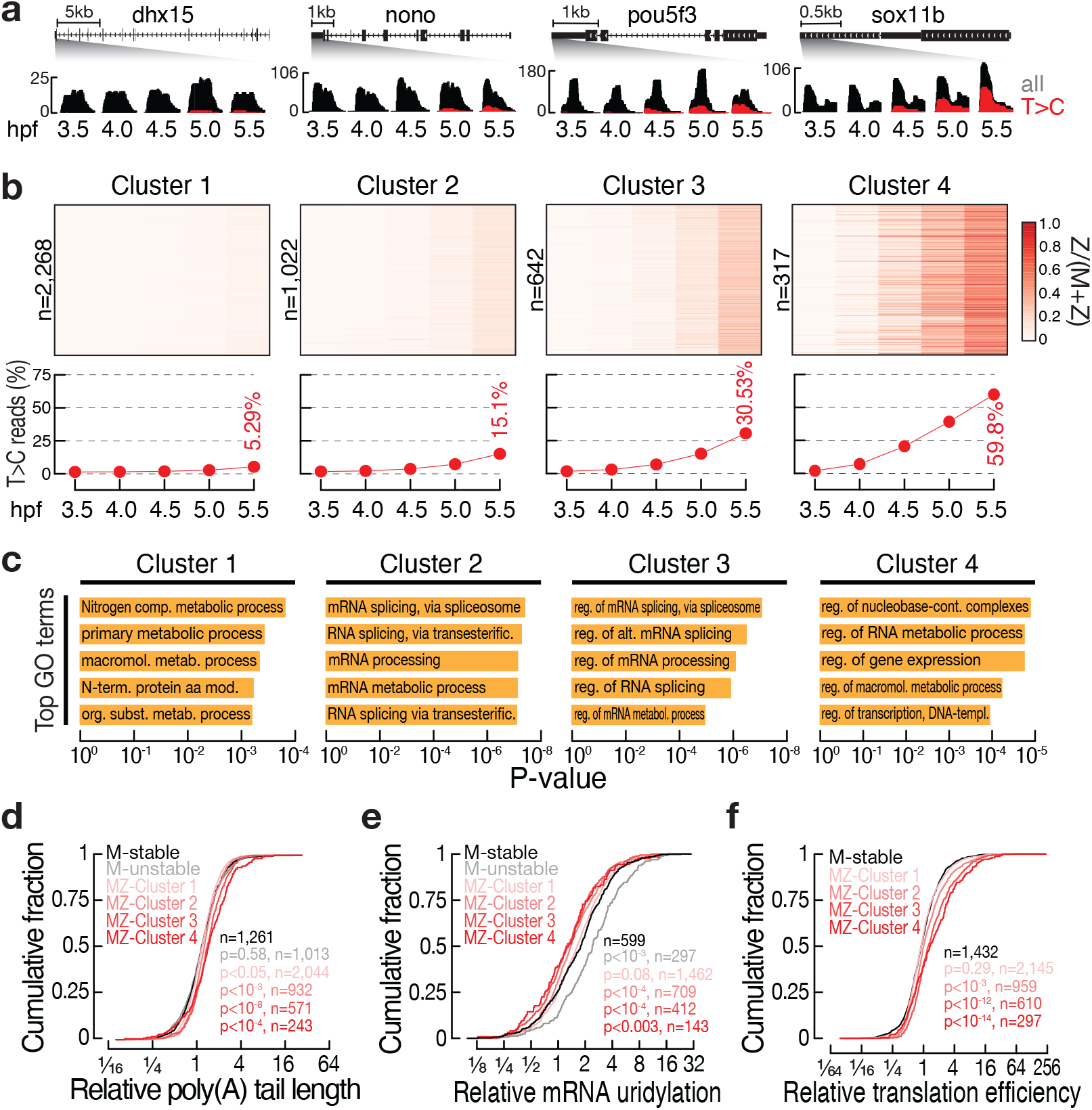
Replacement of MZ transcripts occurs at distinct rates that impinge on signatures of post-transcriptional control. **(a)** Representative genome browser screenshot of MZ genes with constant steady-state expression but distinct rates of zygotic replenishment. Increasing transition kinetics were observed for *DEAH box helicase 15* (*dhx15*), *non POU domain containing octamer-binding* (*nono*), *POU domain class 5 transcription factor 3* (*pou5f3*) and *SRY-box transcription factor 11b* (*sox11b*). Zoom-in to the 3′ end of genes shows steady-state (all, grey) and zygotic expression (T>C, red) as determined by 3′ mRNA SLAMseq at the indicated time points (in hours post fertilization, hpf). **(b)** Heatmap (top) shows the contribution of zygotic transcripts compared to steady-state (Z/[M+Z]) for the indicated number of genes (n) classified into four groups (cluster 1-4) by *k*-medioids clustering of 3′ mRNA SLAMseq at the indicated time points (in hpf). The average contribution of zygotic to steady-state transcripts is shown below and the maximum replacement at 5.5 hpf is indicated. **(c)** Top five gene ontology (GO) terms significantly enriched in each of the gene clusters described in (b). The associated p-value is indicated. **(d** and **e)** Cumulative distribution of the change in poly(A)-tail length (a) and mRNA 3′ end uridylation (b) before and after ZGA (6 vs. 2 hpf). Information on poly(A)-tail length and uridylation was derived from Chang *et al*., 2018 (Chang et al., 2018). The indicated number of genes (n) that we classified into maternal-stable (M-stable; black), maternal-unstable (M-unstable; grey) and MZ clusters (1-4, shades of red; see Fig. 4) are shown. P-values were determined by Komogorov-Smirnov test and are indicated for comparisons of M-unstable with all other transcript categories. **(f)** Cumulative distribution of the change in translation efficiency post- (shield stage, ∼6 hpf) compared to pre-MZT (256 cell stage, ∼2.5 hpf). Translation efficiency was derived from Chew *et al.,* 2013, as determined by ribosome profiling (Chew et al., 2013). The indicated number of genes (n) we classified into maternal-stable (M-stable; black), maternal-unstable (M-unstable; grey) and MZ clusters (1-4, shades of red; see Fig. 4) are shown. P-values were determined by Komogorov-Smirnov test and are indicated for comparisons of M-unstable with all other transcript categories.

GO term analyses revealed differential enrichments for genes of the four MZ clusters with different replacement kinetics: Slowly replenished MZ genes significantly associated with house-keeping functions (cluster 1) or mRNA processing and metabolism (cluster 2). In contrast, genes in cluster 3 and 4 were enriched in gene regulatory functions, including gene networks of protein biosynthesis, mRNA splicing and the proteasome indicating that constant and high mRNA turnover (that must be compensated by active transcription during MZT) (Fig. 5c). Notably, cluster 3 and 4 also include key transcription factors required for ZGA, including *pou5f3* and *nanog*. We concluded that constant steady-state levels of MZ transcripts emerge from balanced synthesis and decay rates that vary substantially between individual genes.

### MZ transition kinetics impinge on hallmarks of post-transcriptional control during MZT

The different rates at which most MZ gene products are replaced by zygotic transcripts presumes the involvement of maternal mRNA clearance mechanisms in the establishment of kinetic transitions. To gain insights into the mechanism(s) that could contribute to the distinct MZ mRNA clearance rates during MZT in zebrafish, we interrogated known features linked to transcript destabilization, including codon adaptation index (CAI), 3′ UTR length, and the presence of miR-430 target sites in maternal and MZ transcripts (Fig. S5d). As expected, M-unstable transcripts were significantly enriched for low CAI and short 3′ UTRs (p<10^-11^, one-sided Fisher’s exact test) and depleted of miR-430 target sites (p<10^-3^), in agreement with previously reported features of rapid maternal mRNA decay (Bazzini et al., 2016; Mishima and Tomari, 2016). Furthermore, M-stable transcripts exhibited significantly higher CAI and longer 3′ UTRs (p<10^-3^), and enrichment for miR-430 targets (p<0.01), also in agreement with their stable nature throughout our time-course and miR-430-mediated decay proceeding thereafter (Fig. S5d) (Mishima and Tomari, 2016; Rabani et al., 2017). While slowly replenished MZ transcripts (cluster 1) mimicked the features observed among M-stable transcripts, i.e., long 3′ UTRs, high CAI and enrichment of miR-430 targets, none of the known features could explain the increasingly more rapid replacement kinetics observed for MZ clusters 2 to 4 (Fig. S5d), implying the involvement of RNA turnover mechanisms that remain to be identified.

To further address how known molecular determinants of mRNA fate and function pertain to the observed rates of MZ replacement, we inspected changes in steady-state poly(A)-tail length, 3′ mRNA uridylation (profiled by Chang *et al*., 2018) and ribosome association (profiled by Chew *et al*., 2013) during MZT. As previously described, poly(A)-tails of maternally deposited mRNAs are short after fertilization and gradually increase during MZT (Chang et al., 2018; Subtelny et al., 2014). While this was generally observed for all classes of genes in our analyses (M-stable, M-unstable and MZ), M-stable transcripts exhibited a significantly stronger increase in poly(A)-tail length compared to M-unstable mRNAs during the first 4 hours (Fig. S5e). This is consistent with cytoplasmic polyadenylation signals that were significantly enriched in M-stable transcripts, but depleted among M-unstable transcripts (Fig. S5f). Following this increase, poly(A)-tail lengths dropped again after 4 hpf, presumably due to the onset of maternal mRNA clearance (Fig. S5e). Interestingly, transcripts of the MZ cluster showed an increase in poly(A)-tail length as rapidly as M-stable mRNAs before ZGA, but retained extended poly(A)-tails for longer thereafter, resulting in a significantly higher poly(A)-tail length for MZ cluster genes compared to M-unstable or M-stable genes (p<0.01, Kolmogorov-Smirnov test) (Fig. 5d). This effect was most pronounced for MZ cluster 4, and correlated with rapid MZ replacement kinetics (Fig. 5b and d). While MZ cluster 1 may accumulate longer poly(A)-tails due to a significant enrichment in cytoplasmic polyadenylation signal (even more so than M-stable), this was not observed for MZ clusters 2 to 4 (Fig. S5f), consistent with the notion that longer poly(A)-tails originate from *de novo* synthesized transcripts (Subtelny et al., 2014). In contrast to polyadenylation, mRNA 3′ uridylation emerges at the onset of MZT and increased in our analyses continuously throughout the time course for both M-stable and M-unstable mRNAs (Fig. S5g) (Chang et al., 2018). In contrast, genes of the MZ cluster appeared to resist mRNA uridylation at the onset of this transition, resulting in a significantly lower penetrance of 3′ uridylation after ZGA when compared to M genes (Fig. 5e). While least apparent for MZ cluster 1, MZ Clusters 2 to 4 showed a significant (p<0.003, Kolmogorov-Smirnov test) and gradually stronger resilience, suggesting that faster MZ replacement kinetics impart increasing resistance against uridylation-mediated mRNA decay (Fig. 5e). A likely explanation for this is that longer poly(A)-tails on *de novo* synthesized transcripts render them less prone to uridylation (Chang et al., 2018). Finally, we investigated the consequences of MZ transition kinetics on translation efficiency (TE). As described before, TE stayed constant for M-stable mRNAs throughout MZT, while M-unstable transcripts dropped in TE quickly after fertilization and remained low thereafter (Fig. S5h). Interestingly, MZ cluster transcripts resembled M-stable transcripts in their TE profiles prior to ZGA, but showed a rapid and significant increase in TE thereafter (p<10^-3^, Kolmogorov-Smirnov test) (Fig. S5h). Again, this effect was gradually stronger for MZ cluster 2, 3, and 4 mRNAs (Fig. 5f and S5h). In summary, our analyses revealed that MZ transition kinetics during MZT impinge on hallmarks of post-transcriptional control that have a direct consequence on ribosome engagement and hence protein output.

## Discussion

Here, we employed metabolic RNA sequencing in zebrafish embryos to resolve the compound (i.e., maternal and zygotic) gene expression during MZT. To this end, we addressed several pre-eminent technical constraints: First, we showed that 3′ mRNA sequencing diminishes biases in the quantification of mRNAs with different poly(A)-tail length (Fig. S1h to l), a crucial feature in light of poly(A)-tail length variants that frequently emerge from cytoplasmic mRNA de- and re-adenylation during zebrafish embryogenesis (Aanes et al., 2011; Winata et al., 2018). Second, we significantly improved on the applicability of 3′ mRNA sequencing to quantify steady-state gene expression (Fig. S1f and g) by systematically re-annotating mRNA 3′ ends (using 3′ GAmES) (Fig. S1d) (Bhat et al., 2021). In fact, we detected between 28,760 and 37,751 mRNA 3′ isoforms across 9 stages of zebrafish embryogenesis (0.75 – 5.5 hpf), among which about two-third are absent from current Ensembl annotations (∼60% intragenic and ∼8% intergenic) (Fig. S1d). And finally, we established an experimental and computational framework for *in vivo* SLAMseq in zebrafish embryos that distinguishes zygotic from maternal transcripts at high sensitivity and accuracy (Fig. 1b-e and Fig.S1 n-o), without perturbing embryonic development (Fig. S1a and b) or normal steady-state gene expression (Fig. S1c). We thereby extend on previous applications of the technology for spatiotemporal mapping of gene expression (Holler et al., 2021). Taken together, we provide a robust experimental framework in zebrafish embryos to accurately measure ZGA in the context of pre-existing maternal mRNAs at unprecedented accuracy, resolution and sensitivity.

By applying 3′ mRNA SLAMseq to zebrafish embryos, our measurements across early zebrafish embryogenesis (up to 5.5 hpf) provided a quantitative view on the overall scope and pattern of ZGA and its contribution to the steady-state transcriptome. In line with previous reports (Heyn et al., 2014), we could ascribe the vast majority (>85%) of robust transcriptional activity immediately after fertilization to the mitochondrial genome. In contrast, nuclear gene expression dominated the *de novo* transcriptome only after 3.5 hpf, thereby demarcating the major wave of ZGA (Fig. 2b-d) (Tadros and Lipshitz, 2009; Vastenhouw et al., 2019). Notably, we detected zygotic expression signals for hundreds of nuclear genes long before the major wave of ZGA and as early as the 2-cell stage (i.e., 0.75 hpf) (Fig. 3a). The underlying genes exhibit hallmark features of early transcribed zygotic genes (Chan et al., 2019; Heyn et al., 2014; Kwasnieski et al., 2019; Pálfy et al., 2020), including few introns, short CDS and signatures of active chromatin such as high accessibility and H3K27 acetylation (Fig. 3d, e and S3d, e). At early timepoints (0.75 hpf and 2 hpf), most of those transcripts could only be detected discontinuously (>79%) and in one of the two replicates (>92%) (Fig. 3b, c and S3b, c), indicative of spurious and/or stochastic expression. Hence, our data significantly prepones and expands the stochastic onset of nuclear gene transcription that was previously observed in a subset of cells and embryos for a much smaller number of purely zygotic genes (Chan et al., 2019; Stapel et al., 2017). Given the reported activity of RNA polymerase II at the multi-copy miR-430 locus as early as the 64-cell stage (Chan et al., 2019), we propose that the widespread spurious transcription at the very initial stages of embryogenesis may be a combined consequence of rapid cell divisions lacking a G2 phase (Heyn et al., 2014; Kwasnieski et al., 2019), stochastic opening of the chromatin, and limiting concentrations of transcription factors (Lee et al., 2013).

At later time points of embryogenesis (i.e., after 3 hpf), nuclear gene transcription is subjected to robust activation that is reinforced gradually rather than in two distinct waves (Fig. 2b, c, S3b), which is in line with previous observations in *Drosophila* and zebrafish (Chan et al., 2019; Heyn et al., 2014; Kwasnieski et al., 2019). Intriguingly, at the blastula stage (i.e., 5.5 hpf), long after the major wave of ZGA, maternal mRNAs still account for >90% of the steady-state transcriptome, a value that is substantially higher than previously estimated (Vastenhouw et al., 2019). Despite the overall dominance of the maternal transcriptome and in line with previous studies (Vastenhouw et al., 2019), we find that most maternal genes (70%) are re-expressed in the zygote, albeit to varying extent (Fig. 4a). In fact, systematic classification based on 3′ mRNA SLAMseq revealed that among 10,342 genes, 14% classify as maternal-unstable (M-unstable; n=1,416), 15% as maternal-stable (M-stable; n= 1569), 67% as maternal/zygotic (MZ; n=6923), and only 4% as purely zygotic (Z; n=434) (Fig. 4a and S4a-d). Notably, most of the zygotic transcriptome (i.e., >84% at the blastula stage) originates from thousands of MZ genes that are on average transcribed at a rate comparable to that of zygotic genes (Fig. 4d and S4e). This is a remarkable finding since research on ZGA has so far almost exclusively focused on the relatively small number of Z genes, mostly because of technical constraints that preempted the unequivocal detection of MZ gene expression. From a biological perspective, the observed gene classes most certainly serve distinct functions, where M-unstable genes may act in reproductive processes, M-stable genes are linked to metabolic and other house-keeping activities, and Z genes fulfil roles in development and differentiation (Fig. 4b). Interestingly, MZ genes were significantly associated with processes related to gene expression – including RNA metabolism, processing and biosynthesis (Fig. 4b) – perhaps indicating a requirement for the re-expression of such genes in sustaining the major wave of zygotic genome activation. Alternatively, we can speculate that replenishment of maternal transcripts that may have been damaged during oogenesis or during the prolonged time of dormancy in the egg, for example due to oxidative damage triggered by reactive oxygen species (Chatzidaki et al., 2021; Murdoch et al., 2001), may be beneficial for embryo survival since this would minimize potential ribosome collisions or cellular stress caused by translation of truncated or erroneous transcripts.

The unexpectedly broad scope of maternal gene activity during MZT raises important open questions in the regulation and dynamics of MZ genes in the embryo. In fact, *k*-medioids clustering revealed four distinct groups of MZ genes that exhibit constant steady-state expression during MZT but differ substantially in the rates at which maternal transcripts are replaced by zygotic gene products (Fig. 4a and S4a), resulting in a gradually increasing median zygotic contribution of up to 5.3% (for cluster 1) to up to 59.8% (for cluster 4). Slowly replenished MZ genes significantly associated with house-keeping functions (cluster 1) or mRNA processing and metabolism (cluster 2) perhaps reflecting evolutionary adaptation of gene expression kinetics to energy constraints, as proposed before (Herzog et al., 2017a; Schwanhäusser et al., 2011). In contrast, genes in cluster 3 and 4 were enriched in gene regulatory functions, including gene networks of protein biosynthesis, mRNA splicing and the proteasome indicating that constant and high mRNA turnover (that is compensated by the activation of transcription during MZT) may allow for rapid adaptation to changing developmental and/or environmental changes (Fig. 5c). Intriguingly, none of the known features linked to maternal mRNA clearance during MZT in zebrafish (i.e., codon-adaptation index, 3′ UTR length, and the presence of miR-430 binding sites) could explain rapid MZ transition kinetics (Fig. S5d), highlighting a new avenue of future research to identify the underlying mechanism(s). Irrespective of the precise mode of action, such different kinetics may be a consequence of inherent and conserved gene features (Fig. S5c), with immediate implications for the persistence of genetic information provoked by the specific functions of the encoded proteins and their regulation.

At the molecular level, MZ replacement kinetics alter the hallmarks of post-transcriptional control and protein output. We found that the fastest replaced cluster (cluster 4), which is enriched for transcripts encoding proteins involved in the regulation of RNA expression and processing, showed features that are consistent with a demand for the efficient translation of such transcripts in the embryo: significantly elevated poly(A)-tail length, low U-tailing, and high TE (Fig. 5d-f and S5). We therefore provide evidence that rapid replenishment of those transcripts contributes to the efficient *de novo* protein production in the embryo to enable robust progression through embryogenesis. Consistent with this view, key transcription factors required for ZGA, including Pou5f3 and Nanog, are encoded by rapidly replenished MZ genes.

## Supplemental Information

Supplemental information includes five Supplemental Figures (S1-S5).

## Acknowledgements

We thank Eivind Valen and all members of the Ameres and Pauli laboratories for technical support and helpful discussions. RNA sequencing was performed at the VBCF NGS Unit (www.vbcf.ac.at). This research was supported by the Austrian Science Fund FWF (SFB-F80 to SLA and AP and FWF START program Y1031-B28 to AP), the European Research Council (ERC-CoG-866166 RiboTrace and ERC-PoC-825710 to SLA), the Austrian Research Promotion Agency (Headquarter grant FFG-852936 to AP) the Human Frontier Science Program (HFSP) Career Development Award (CDA00066/2015 to AP), the Austrian Academy of Sciences, Boehringer Ingelheim and a Boehringer Ingelheim Fonds (BIF) PhD fellowship to LECQ.

## Author Contributions

AP, SLA and PB conceived the study. PB conducted bioinformatic analyses. LECQ performed zebrafish experiments and 3′ mRNA SLAMseq libraries. VAH and NF performed initial SLAMseq experiments. SLA and AP supervised the study. PB, AP and SLA wrote the manuscript with input from all authors.

## Declaration of Interests

VAH and SLA declare competing interest based on a granted patent related to SLAMseq. SLA is co-founder, advisor, and member of the board of QUANTRO Therapeutics GmbH.

## Supplementary Figure Legends

**Figure S1. Time-resolved mRNA 3**ʹ **end sequencing in zebrafish embryos distinguishes zygotic from maternally deposited mRNA.** Referring to Fig. 1. **(a)** Phase-contrast images at the indicated time points (in hours post fertilization, hpf) of untreated zebrafish embryos and embryos injected at the one-cell stage with the indicated concentrations of 4-thiouridine (4sU in mM). **(b)** Quantification of developmental progression of the experiment shown in (a). Embryos were categorized as ‘normal’, ‘abnormal’, and ‘dead’ based on phenotypic appearance for the indicated experimental condition and developmental stage. The total number of inspected embryos is shown (n). **(c)** Comparison of steady-state gene expression (in reads per million, rpm) at the indicated time points (in hpf) of untreated zebrafish embryos (–4sU) and embryos injected with 1.5 mM 4sU (+4sU) as determined by 3′ mRNA sequencing. The number of genes (n; rpm>1), Pearson correlation coefficient (R_P_) and associated p-value (p) is indicated. **(d)** Overview of the 3′GAmES pipeline used for the systematic annotation of mRNA 3′ ends in zebrafish, as previously described(Bhat et al., 2021). Processing seps (grey) and required input files (black) are indicated. Note, that mRNA 3′ ends from each developmental stage were retained only if they were identified in at least two consecutive developmental stages. **(e)** Statistics of 3′ GAmES input and output. **(f)** Impact of 3′GAmES-derived annotations on the quantitative representation of transcripts in 3′ mRNA sequencing datasets compared to conventional RNA sequencing approaches. mRNA sequencing (RNAseq in transcripts per million, tpm) and 3′ mRNA sequencing (3′ mRNAseq in reads per million, rpm) in 0.75 hpf zebrafish embryos was compared, relying on Ensembl- (left) or 3′GAmES-derived annotations (right). Number of comparisons (n) were restricted to genes that exhibited > 5 tpm in RNAseq data. Spearman correlation coefficients (R) and associated *p*-values (p) are indicated. **(g)** Overview of correlation analysis as described in (f) performed in zebrafish embryos at the indicated time points (in hpf). Spearman correlation coefficients (R) are indicated. **(h, k)** Contour plots report relative (rel.) transcript abundance comparing 3′ mRNAseq (top, green) or poly(A)-selected (poly[A]^+^) RNAseq (bottom, brown) with rRNA-depleted (ribo^−^) RNAseq across mRNAs with defined poly(A) tail lengths (in nucleotides, nt) as described previously by (Subtelny et al., 2014) (h) or (Chang et al., 2018) (k). **(i, l)** Boxplot representation of analyses shown in (h) and (k), respectively. Data was binned according to even-sized poly(A) tail length quantiles and statistically significant (multiple comparison-corrected Wilcoxon test) over or underrepresentation in expression ratios is indicated in green (3′ mRNAseq vs. ribo– RNAseq, top) or brown (polyA+ RNAseq vs. ribo– RNAseq, bottom), while non-significant boxplots are colored in grey. **(m)** Normalized steady-state expression of pre-defined zygotic (left) high-confidence (h.-c.) maternal and suspected (susp.) maternal genes at the indicated time points (in hpf). Expression was assessed by 3′ mRNA sequencing. Each gene is reported relative to the expression (in ppm) at which it is highest abundant. The number of genes for each group is indicated (n). **(n)** T>C conversion (conv.) rate for suspected maternal transcripts (see main text) that exhibited a zygotic-like or maternal-like accumulation of T>C conversion containing reads (Fig. 1 c and main text) in 3′ mRNA SLAMseq datasets derived from zebrafish embryos at the indicated developmental stage (in hpf). Individual genes for which a confident average conversion rate could be derived from two independent biological replicates (n = number of genes) are shown. P-values (Mann-Whitney test; p) are indicated. **(o)** Steady-state abundance (ppm) and T>C conversion rate (%) at the indicated time points (in hpf) is shown for three example genes of suspected maternal genes (see Fig. 1c and d) that exhibit zygotic contribution in 3′ mRNA SLAMseq datasets. **(p)** T>C conversion (conv.) rate for pre-defined zygotic transcripts (see main text) in 3′ mRNA sequencing datasets derived from untreated zebrafish embryos at the indicated time point (in hpf). Individual genes for which a confident average conversion rate could be derived from two independent biological replicates (n = number of genes) are shown. P-value (Mann-Whitney test; p) is indicated. **(q)** Comparison of steady-state gene expression (in reads per million, rpm; left) or de novo transcription (T>C reads per million, rpm; right) between two independent replicates of 3′ mRNA SLAMseq experiments at the indicated time point (hpf). Only genes expressed at >1 (T>C) read per million (rpm) in both replicates were considered. The number of comparisons (n) is shown. Pearson correlation coefficients (R_P_) and associated p-values are presented.

**Figure S2. The transcriptional landscape of zebrafish embryogenesis.** Referring to Fig. 2. **(a)** Contour plots show a comparison of *de novo* transcription (i.e., T>C conversion containing reads in rpm, red) or background nucleotide conversion errors (T>A conversion containing reads in rpm, black) with steady-state gene expression (in rpm) in zebrafish embryos injected with 4-thiouridine (+4sU, red) or in untreated control embryos (−4sU, black) as determined by 3′ mRNA SLAMseq at the indicated time point (in hours post fertilization, hpf). Boxplots show a quantitative comparison of *de novo* transcription signal or background conversions by comparing T>C and T>A conversion containing reads (in rpm). P-value was determined by Wilcoxon test. The number of inspected genes is indicated (n). **(b)** Boxplots for pre-defined zygotic and high-confidence (h.-c.) maternal genes (see also Fig. 1c and d) report the fraction of reads containing the indicated number of T>C conversion per read in 3′ mRNA SLAMseq data generated from zebrafish embryos at 5.5 hpf.

**Figure S3. Evidence for the spurious transcription at hundreds of gene loci prior to MZT.** Referring to Fig. 3. **(a)** True and false positive rates for the detection of newly synthesized transcripts in 3′mRNA SLAMseq data generated from zebrafish embryos at 5.5 hpf. The discovery rate for pre-defined zygotic (n=47; true positive) and high-confidence maternal genes (n=25; maternal) depending on T>C confidence score (PTC; see Fig. 2e and main text) is shown for two independent biological replicates. The PTC threshold used to identify genes with transcriptional signal at early embryogenesis is indicated (dashed line). **(b)** Venn diagrams report the overlap of genes detected above PTC-cutoff (see a) in two independent biological 3′ mRNA SLAMseq replicates at the indicated time of embryo development (hpf). Significance of overlap was determined by Fisher’s exact test. **** = p<10^-4^. **(c)** Heatmap for all genes detected above PTC-cutoff at 2 hpf (n=554) reports the presence of T>C conversion containing reads (above PTC-cutoff) in none (white), one (grey) or both (black) of two independent 3′ mRNA SLAMseq experiments at the indicated subsequent time points (in hpf). Genes with continuous (red) or discontinuous signal (blue) in at least one of two replicates throughout the entire time course are highlighted (left). **(d)** Median number of introns (left) or length of the coding sequence (CDS in kilobases, kb) for the indicated number of genes ranked by highest (filled circles) or lowest (open circles) T>C confidence score (PTC) in 3′ mRNA SLAMseq datasets from zebrafish embryos at 2 hpf. Statistically significant deviation (Wilcoxon test; ****=p<10^-4^) from median (dashed line) and interquartile range (grey area) of all genes in the dataset is indicated. **(e)** Normalized enrichment scores (NES) of ATACseq (left) and H3K27 acetylation (right) for the indicated number of genes ranked by highest (red) or lowest (blue) T>C confidence score (PTC) in 3′ mRNA SLAMseq datasets generated from zebrafish embryos at 2 hpf. Significant enrichment (hypergeometric test with Benjamini-Hochberg correction; p<0.05) is indicated in opaque colors.

**Figure S4. Classification of the zebrafish transcriptome during MZT.** Referring to Fig. 4. **(a)** Schematic overview of the criteria used to classify the zebrafish embryonic transcriptome based on 3′ mRNA SLAMseq datasets. Parameters used to group genes into maternal-unstable (black), maternal-stable (blue), maternal-zygotic (yellow) and zygotic (red) are shown. **(b)** Correlation between steady-state gene expression of the indicated number of genes (n) classified into maternal-unstable (M-unstable; black), maternal-stable (M-stable; blue), maternal-zygotic (MZ; yellow) and zygotic (Z; red) in fertilized oocytes and 2-cell stage embryos (0.75 hpf). Pearson correlation coefficient (R) and associated p-value (p) are indicated. **(c)** Pie-charts show the fraction of zygotically expressed genes reported by Heyn *et al*., 2014 (left), Harvey *et al*., 2013 (middle) and Lee *et al*., 2013 (right) that were classified by 3′ mRNA SLAMseq as maternal-unstable (M-unstable; dark grey), maternal-stable (M-stable; blue), maternal-zygotic (MZ; yellow), zygotic (Z; red) and undetected (light grey). The total number of analyzed genes is indicated (n). Note, that only genes with consistent annotation in the original study and the zebrafish genome assembly GRCz11 were considered. **(d)** T>C conversion (conv.) rate for MZ or Z genes newly classified in this study. Time points are indicated (in hpf). The number of analyzed genes is shown (n). **(e)** Quantification of zygotic expression (i.e., T>C conversion containing reads in rpm) per gene for zygotic (Z; red) or maternal-zygotic (MZ; yellow) genes (as classified in Fig. 3a) at the indicated time points (in hpf). Median (filled circles) and interquartile range (error bars) is shown. The number of genes is indicated (n). The relative (rel.) median transcriptional (trx.) output per gene reports the relative median T>C conversion containing reads detected per zygotic (red) or maternal-zygotic (yellow) gene at the indicated time point.

**Figure S5. Replacement of MZ transcripts occurs at distinct rates that impinge on signatures of post-transcriptional control.** Referring to Fig. 5. **(a)** View of Fig. 5b across the entire time course. The average contribution of zygotic to steady-state transcripts for the indicated number of genes (n) belonging to the MZ clusters defined in Fig. 5 is shown. **(b)** Steady-state abundance at 0.75 hpf (in reads per million, rpm) of transcripts belonging to the indicated MZ cluster (shades of red) and of all detected genes (black). The number of analyzed genes is indicated (n). P-values based on Wilcoxon test are reported. **(c)** Half-lives of murine transcripts that are orthologous to the indicated zebrafish MZ clusters (shades of red) and of all detected genes (black) in mouse embryonic stem cells (mESC). The number of analyzed genes is indicated (n). P-values based on Wilcoxon test are reported. **(d)** Scatterplots show codon adaptation index (COI) and 3′ UTR length (in nucleotides, nt) for the indicated genes classified into maternal-stable (M-stable), maternal-unstable (M-unstable) and MZ clusters (1-4; see also Fig. 4). The median of 3′ UTR length (628 nt) and CAI (0.807) of all transcripts were used for thresholding (dashed lines). Percentages of genes falling in each quadrant are indicated. Percentages of genes falling in each quadrant are indicated and highlighted in case of significant enrichment (red) or depletion (blue). Quadrants in which genes exhibit a significant enrichment (red boxes) or depletion (blue boxes) of miR-430 target sites are highlighted. Enrichments and depletions are evaluated for significance using the Fisher’s exact test (p-value cutoff at p<0.01). **(e)** Average poly(A)-tail length (in nucleotides; nt; a) for mRNAs classified into maternal-stable (M-stable; black), maternal-unstable (M-unstable; grey) and MZ clusters (1-4, shades of red; see also Fig. 4) at the indicated time point (in hpf). Information on poly(A)-tail length was derived from Chang *et al*., 2018 (Chang et al., 2018). **(f)** Odds ratio (Fisher’s exact test, significance highlighted in black if p<0.01) for the enrichment (>1) or depletion (<1) of previously identified targets of cytoplasmic polyadenylation in the indicated group of genes (Winata et al., 2018). **(g)** Average fraction of 3′ uridylation for mRNAs classified into maternal-stable (M-stable; black), maternal-unstable (M-unstable; grey) and MZ clusters (1-4, shades of red; see also Fig. 4) at the indicated time points (in hpf). Information on mRNA 3′ uridylation was derived from Chang *et al*., 2018 (Chang et al., 2018). **(h)** Translation efficiency for mRNAs classified into maternal-stable (M-stable; black), maternal-unstable (M-unstable; grey) and MZ clusters (1-4, shades of red; see also Fig. 5) at the indicated time points (in hpf). Information on translation efficiency was derived from Chew *et al*., 2013 (Chew et al., 2013).

## RESOURCE AVAILABILITY

### Contact for reagent and resource sharing

Further information and requests for resources and reagents should be directed to and will be fulfilled by Stefan L. Ameres and Andrea Pauli.

### Data and code availability

All data generated for this study have been deposited at GEO (GSE185283) and will be publicly available as of the date of publication.

The scripts used for the analysis presented in this study are available on GitHub (https://github.com/AmeresLab/zebrafishAnalysis).

## EXPERIMENTAL MODEL AND SUBJECT DETAILS

Zebrafish (*Danio rerio*) were raised according to standard protocols (28°C water temperature; 14 h light/10 h dark cycle). TLAB fish were generated by crossing zebrafish AB and the natural variant TL (Tupfel Longfin) stocks, and served as wild-types for all experiments. All zebrafish experiments were conducted according to Austrian and European guidelines for animal research and approved by local Austrian authorities (animal protocols GZ: 342445/2016/12 and MA 58-221180-2021-16).

## METHOD DETAILS

### Metabolic RNA labeling with 4sU

Wild-type (TLAB) embryos (derived from mating of an individual male-female pair; two independent biological replicates) were collected within 15 min of mating. Embryos were dechorionated (1 mg/ml Pronase) and within 30 min after egg-laying injected with 4sU (1.5 mM [unless indicated otherwise] final concentration in the embryo, corresponding to the injection of 1 nl of a 45 mM 4sU injection mix into the cell of a 1-cell stage embryo). Injected embryos were kept in six-well agarose-coated plates in E3 medium (5 mM NaCl, 0.17 mM KCl, 0.33 mM CaCl_2_, 0.33 mM MgSO_4_, 0.1% [w/v] Methylene Blue) at 28°C in the dark until further processing (see 3′ mRNA SLAMseq). Samples were only used for RNA isolation if the rate of fertilization was higher than 60%. Fertilization rates were quantified after 3 hours as the rate of embryos showing normal cell cleavage. For +4sU time-courses, two biological replicates were collected, each using a single (different) pair of parents. In parallel, control samples without 4sU injection were collected at the indicated time points.

For toxicity measurements, injected embryos were inspected under a stereomicroscope at the indicated time points (1-24 hpf), and changes in morphology were quantified manually. Pictures of embryo development were taken on a ZEISS Stemi 508 stereo microscope with camera (2x magnification, FlyCapture2 software).

### Total RNA isolation

Per stage and replicate, 30 zebrafish embryos were homogenized in 500 µl of TRIzol (Invitrogen) and total RNA was extracted according to the instructions of the manufacturer. During precipitation and washes of +4sU samples, 0.1 mM DTT was supplemented, as previously described (Herzog et al., 2017b). RNA quality was assessed on a capillary electrophoresis instrument (AATI Fragment Analyzer) and quantified on a microvolume spectrophotometer (Nanodrop).

### 3′ mRNA SLAMseq

Total RNA was processed according to the standard SLAMseq protocol, described previously (Herzog et al., 2017b, 2017a). Briefly, 5 μg of total RNA was treated with 10 mM iodoacetamide in alkylation buffer (50 mM sodiumphosphate buffer, pH8; 50% DMSO) at 50°C for 15 min, followed by ethanol precipitation. For each sample, 500 ng of total RNA were used as input for QuantSeq (QuantSeq 3′ mRNA-Seq Library Prep Kit FWD for Illumina; Lexogen) following the autoQuantSeq automated workflow. Libraries were assessed for quality on a capillary electrophoresis instrument (AATI Fragment Analyzer), multiplexed to equimolar concentrations, and sequenced on a HiSeq 2500 system (Illumina) on SR100 mode.

### Whole-transcriptome RNA sequencing

For whole transcriptome RNA library preparation, zebrafish embryos were collected at the indicated developmental time points and processed as described above. Enrichment of polyadenylated RNAs and depletion of ribosomal RNA from total RNA samples was performed from 1 µg of total RNA using the Poly(A) RNA Selection Kit (Lexogen) or RiboCop rRNA Depletion Kit (Lexogen), respectively, according to the instructions of the manufacturer. Stranded cDNA libraries were generated using NEBNext Ultra Direction RNA Library Prep Kit for Illumina (New England Biolabs) and indexed with NEBNext Multiplex Oligos for Illumina (Dual Index Primer Set I; New England Biolabs). Libraries were assessed for quality on a capillary electrophoresis instrument (AATI Fragment Analyzer) and sequenced on a HiSeq 2500 instrument (Illumina) in PE-50 mode.

### Data analyses

#### Custom mRNA 3′ end annotation

For each developmental time point, mRNA 3′ ends were defined using 3′GAmES, as reported previously (Bhat et al., 2021). To combine the annotations from different stages, only high confidence ends (+/- 5nts) detected in at least two consecutive stages were considered. A final set of counting windows (250 nt in length) were created by combining 3′GAmES-derived high confidence ends and ENSEMBL 3’ UTR annotations from GRCz11. Overlapping counting windows were merged.

#### Selection of h.-c. maternal susp. maternal and zygotic gene sets

High-confidence (h.-c.) M genes, suspected (susp.) M genes, and Z genes (described in Fig. 1c and d) were identified by combining rRNA-depleted whole transcriptome sequencing and 3′ mRNA sequencing as well as previously published classifications. High-confidence. M genes were derived from genes classified as maternal by Lee *et al*., 2012 and further refined based on the following criteria: (1) >1 rpm in 3′ mRNA sequencing datasets at 0.75 hpf; (2) >5 tpm at 2-cell stage and (tpm at 28 hpf)/(tpm at 2hpf) < 0.1 in whole transcriptome RNAseq datasets from Pauli *et al*., 2021; (3) association with GO terms ‘oogenesis’, ‘meiosis I’, ‘meiosis II’, but not ‘mitosis’ in zebrafish (Carbon et al., 2008). Suspected M genes were defined as reported by Lee *et al*., 2012 and filtered based on (rpm at 4.5 hpf)/(rpm at 1 hpf) < 0.5 in whole transcriptome RNAseq datasets generated in this study. Z genes were defined by combining zygotic genes reported by Heyn *et al*., 2014, Harvey *et al*., 2013, and Lee *et al*., 2014 and filtered according to steady-state expression in 3′ mRNA seq datasets ([rpm at 0.75 hpf] = 0, and [rpm at 5.5 hpf] > 0).

#### Analyses of 3′ mRNA SLAMseq datasets

Adapters were trimmed using cutadapt (version 1.18) and reads shorter than 17 nucleotides after trimming were discarded (Martin, 2011). SLAMdunk (version 0.3.4) was used for further processing, mapping, filtering and counting T>C conversions (Neumann et al., 2019). Specific parameters were as follows: Adenine stretches with a length of >4 nucleotides at the 3′ end of reads were trimmed (-a 5); a minimum identity of 95% was required during mapping (-mi 0.95); multimapping was allowed for a maximum of 100 times (-n 100); and a minimum variant fraction (i.e., nucleotide conversion rate at a specific position) of 0.2 was used (-mv 0.2) for SNP masking. More precisely, T>C conversions were evaluated only after accounting for genotype specific SNPs. If a conversion was present at the same genomic position in >20% of reads of any given library or cumulatively across all libraries of a time-course from the same replicate, the conversion was considered as a SNP and masked from further analyses. To this end, reads from mapped files were piled up using samtools/1.4-foss-2017a (base quality computation disabled) (Li et al., 2009). SNPs were called using VarScan.v2.4.1 on the combined bam files (Koboldt et al., 2012), positions covered by more than >10 reads were retained (--min-coverage 10), SNPs were removed based on a per position threshold of 20% (--min-var-freq 0.2), and strand filtering was set to 0 (--strand-filter 0). SNP masking was also extended to T>A conversions in the cases where background sequencing errors were determined. For all conversions only positions with a base quality of >26 were retained. Conversion rates were calculated as the ratio of number of T>Cs converted by the number of Ts covered for each counting window. Background error rates were estimated based on linear regression of T>A conversions (in sample control). Background error rates were subtracted from T>C conversion rates prior to any of the downstream analyses.

#### Mapping of whole transcriptome RNA sequencing datasets

Whole transcriptome RNA sequencing data was mapped to the zebrafish genome (GRCz11) using STAR aligner (star/2.5.2b- foss-2017a), whereby unique mapping was performed with 2-pass filtering (Dobin et al., 2012). Reads were counted using feature counts from the package subread/1.5.0-p1-foss-2017a (Liao et al., 2013, 2014). Browser tracks were created using bedtools (v2.27.1) and the ucsc-kent-utils package (Kent et al., 2010; Quinlan, 2014).

#### Classification of the MZT transcriptome

The transcriptome was classified based on normalized gene expression and T>C conversion rates across the entire time course. All counting windows with expression >5RPM at any time point were considered for the analysis. First, *K*-medoids clustering was performed on normalized gene expression over time using the R package cluster (Maechler et al., 2022). Clustering resulted in 3 clusters with (i) decreasing gene expression, (ii) constant gene expression, (iii) increasing gene expression. These 3 groups were further inspected by *K*-medioids clustering according to T>C conversion rates, resulting in the following sub-clusters: Cluster (i) resulted in three sub-clusters with no accumulation of T>C conversions (one sub-cluster; assigned to M), or increasing T>C conversion rates (two sub-clusters; assigned to MZ). Cluster (ii) also separated into three subclusters with no T>C conversions (two sub-clusters, assigned to M-stable) or increasing T>C conversion rates (one sub-cluster; assigned to MZ). Lastly, genes of Cluster (iii) were further analyzed based on expression at 0.75 hpf and 2 hpf: Genes detected at <1 rpm at both time-points and increasing steady-state expression as well as T>C conversions were classified as Z; and genes detected at ≥ 1 rpm at both time-points and increasing steady-state expression as well as T>C conversion rates were assigned to MZ. A summary for the classification strategy is provided in Fig. S4a.

#### Gene ontology (GO) term analyses

GO term analysis on gene subsets was performed using GOrilla (Eden et al., 2009).

#### Processing of external datasets

CAGE data was derived from Haberle *et al*., 2014 (NCBI Sequence Read Archive: SRA097279) and mapped to the zebrafish genome (GRCz11) using bowtie2 (v2.2.9) (Langmead and Salzberg, 2012). BigWig files were created using bedtools.v.2.27.1 (Quinlan, 2014).

ATAC-seq data was downloaded as BigWig files from Pálfy *et al*., 2020. ATACseq signal was determined within a 1kb window upstream and downstream of the transcription start site for each gene.

H3K27Ac ChIPseq data were derived from Bogdanović et al., 2012. FastQ files were downloaded, adapter-trimmed using cutadapt (1.18), and mapped using bowtie2 (2.3.5.1) (Langmead and Salzberg, 2012; Martin, 2011). Coverage was determined within 1kb windows upstream and downstream of the transcription start site of each gene using bedtools (2.27.1)(Quinlan, 2014). The coverage values were used for gene set enrichment analyses (using GSEA v4.0.1) to check for enrichment of signal (Subramanian et al., 2005). Binding sites of miR-430 were identified using TargetScan6.2 (Ulitsky et al., 2012), and applied to 3′UTR sequences derived from ENSEMBL annotations adjusted to mRNA 3′ end annotation as described above. Genes responsive to miR-430 inhibition were identified based on previously published data (Mishima and Tomari, 2016). Briefly, datasets of RNAseq experiments at 6 hpf (± anti-miR-430 morpholino treatment) were downloaded (GEO accession GSE71609) (Mishima and Tomari, 2016), and mapped to the zebrafish genome (GRCz11) using STAR v.2.5.1 (Dobin et al., 2012). Reads were counted using featureCounts from the Subread package v2.0.0 (Liao et al., 2014). Differential expression analysis was performed using DESeq2 (Love et al., 2014). Targets of miR-430 were identified based on the presence of a target site in the 3’ UTR (TargetScan prediction) and at least 1.5-fold upregulation upon miR-430 morpholino-treatment.

TAIL-seq data was derived from Chang *et al*., 2018 and downloaded from the supplementary data. Poly(A)-tail length and uridylation status was downloaded and processed in R v.3.5. The weighted average of poly(A)-tail lengths was calculated per gene to quantify poly(A) tail lengths.

Ribosome profiling data and RNA-seq data were obtained from Pauli *et al*., 2012 (GEO accession number GSE32898) and Chew *et al*. 2013 (GEO accession number GSE46512). Trimming of the adaptors was performed using Cutadapt v1.12.0 (Martin, 2011), and reads shorter than 20 nucleotides after trimming were discarded. For ribosome profiling and RNA-seq, trimmed reads were mapped against the rRNA sequences from the SILVA rRNA database using Bowtie 2 (-N 1 -L15) (Langmead and Salzberg, 2012; Quast et al., 2013). Reads that matched rRNA were not considered for downstream analyses. The remaining ribosome-protected fragments (RPF) or RNA-seq reads were mapped against the zebrafish genome (GRCz10) with Tophat2 (-g 1 -x 1) aided with the gene models from ENSEMBL version 81 (Cunningham et al., 2014; Trapnell et al., 2009). Mapped reads were quantified using HTSeq (Putri et al., 2022). For RNA-seq analyses, full-length transcripts were considered while for RPF only coding sequences were taken into account. Read counts were converted to FPKM, where FPKM = [(read counts) / (length × total read counts)] × 10^9^. To calculate translational efficiencies (TE), RPF (in FPKM) were divided by RNAseq signal (in FPKM).

Codon adaptation index (CAI) was used as determined previously (Cabrera-Quio et al., 2021). Median CAI was found to be 0.806. For each counting window, the most proximal annotated 3′ UTR start (Annotated by ENSEMBL and refSeq GRCz11 annotations) was used (Cunningham et al., 2014; O’Leary et al., 2016). Target sites of miR-430 were predicted in the respective 3′ UTRs. For each gene, the average 3′ UTR lengths of all isoforms was determined, and the median 3′ UTR lengths of all genes in the annotation was calculated to be 628nts.

## QUANTIFICATION AND STATISTICAL ANALYSIS

All software used in this study is referred to and cited in the method details section. The statistical details of experiments including statistical tests used, number of replicates and the number of data points used (n) are denoted in the relevant figures and in the main text. All statistical tests, and correlations were analyzed using R v.3.5. P-values were adjusted to multiple comparisons as described. A p-value of < 0.05 was considered statistically significant unless indicated otherwise.

