## Supplementary material for "SLAMseq resolves the kinetics of maternal and zygotic gene expression in early zebrafish embryogenesis": Figure S1

Bhat et. al., Figure S1

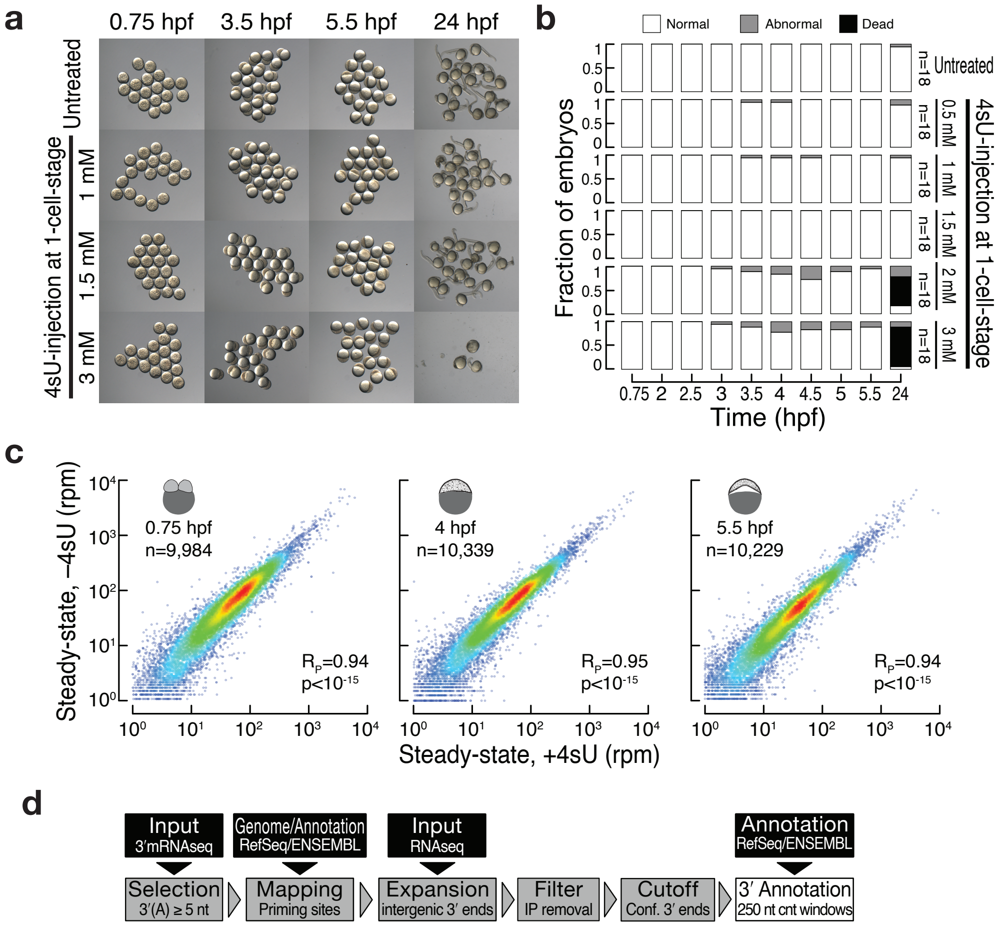


e

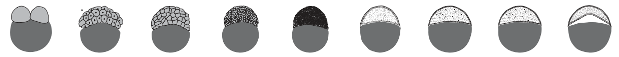

| hpf: | 0.75 | 2 | 2.5 | 3 | 3.5 | 4 | 4.5 | 5 | 5.5 |
|---|---|---|---|---|---|---|---|---|---|
| Input reads (mio): | 40.2 | 19.9 | 38.2 | 26.5 | 29.2 | 28.7 | 20.9 | 20.5 | 29.7 |
| expr. genes (>=2rpm): | 9378 | 9070 | 9433 | 9524 | 9470 | 9726 | 9190 | 9264 | 9083 |
| **3´isoforms** | | | | | | | | | |
| all: | 37751 | 28760 | 35029 | 32177 | 34916 | 34833 | 31590 | 34373 | 31952 |
| annotated: | 11040 | 9967 | 11042 | 10943 | 10902 | 11152 | 10301 | 10774 | 10515 |
| new intragenic: | 23265 | 16086 | 20600 | 18191 | 20779 | 20617 | 19380 | 21573 | 19582 |
| new intergenic: | 3446 | 2707 | 3387 | 3043 | 3235 | 3064 | 1909 | 2026 | 1855 |

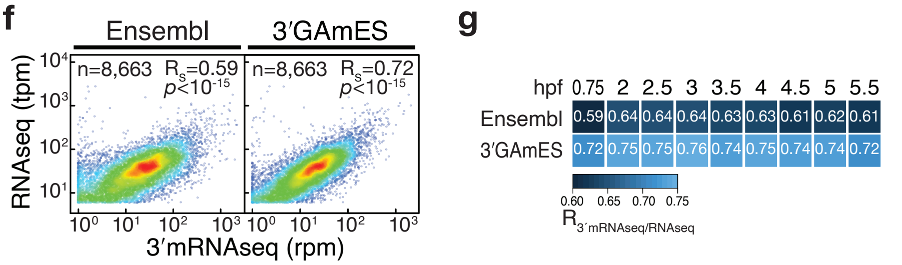

Bhat et. al., Figure S1

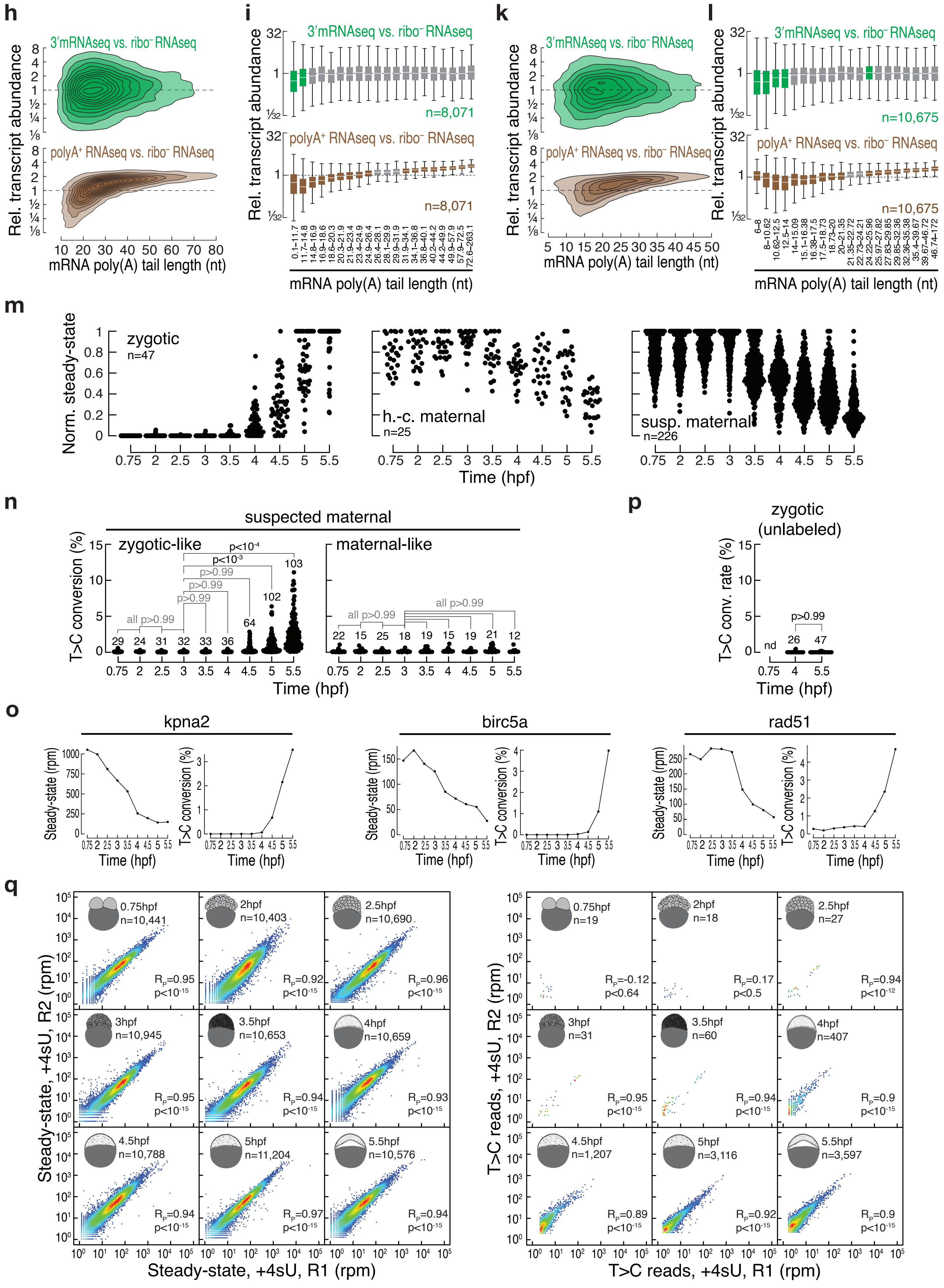
