## Supplementary figures and images for "SLAMseq resolves the kinetics of maternal and zygotic gene expression in early zebrafish embryogenesis"

### Figure S2

Bhat et. al., Figure S2

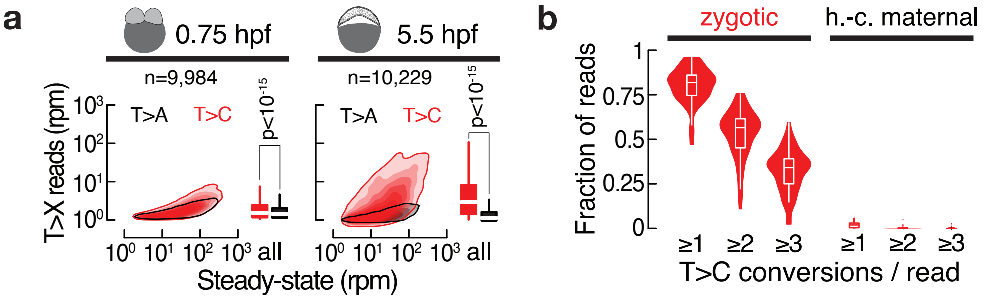

### Figure S3

Bhat et. al., Figure S3

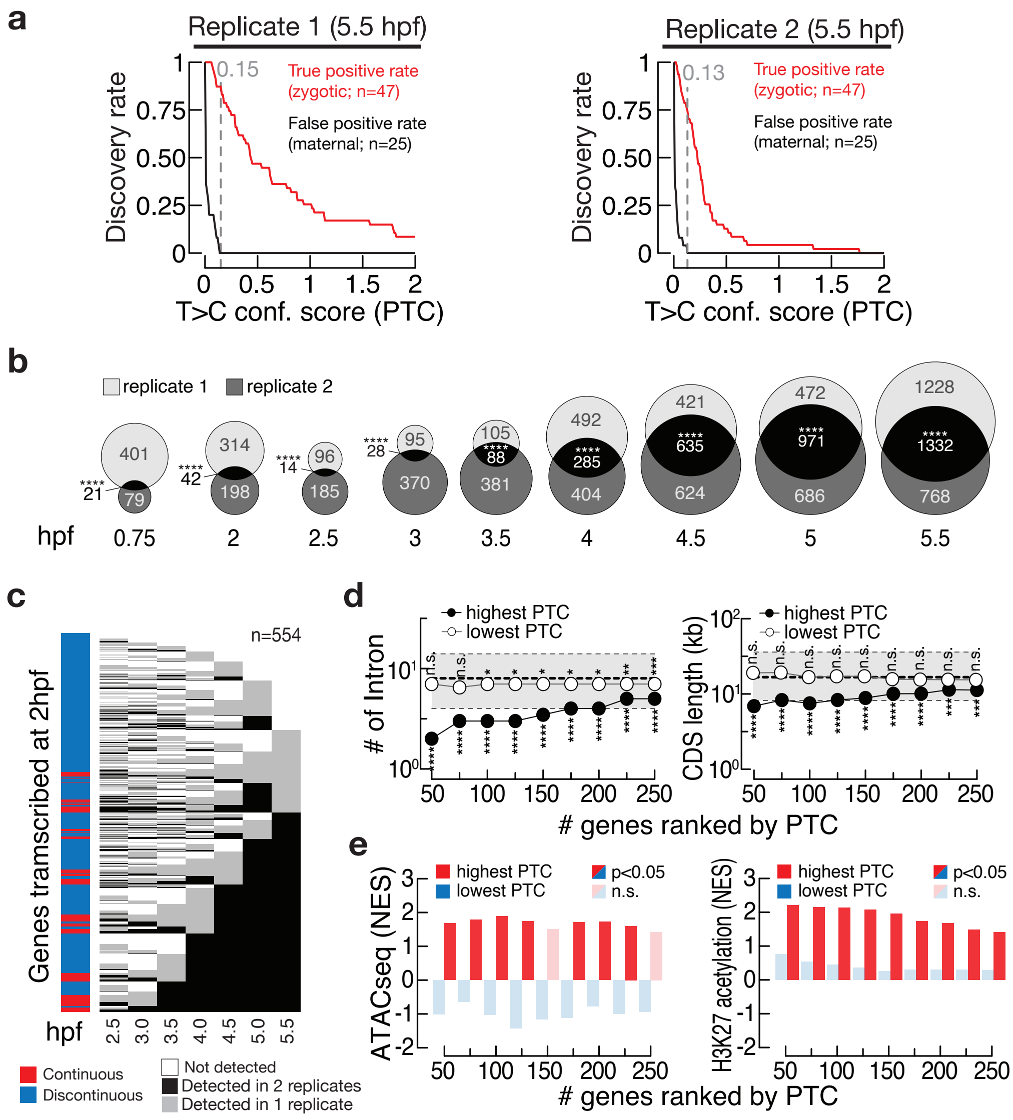

### Figure S4

Bhat et. al., Figure S4

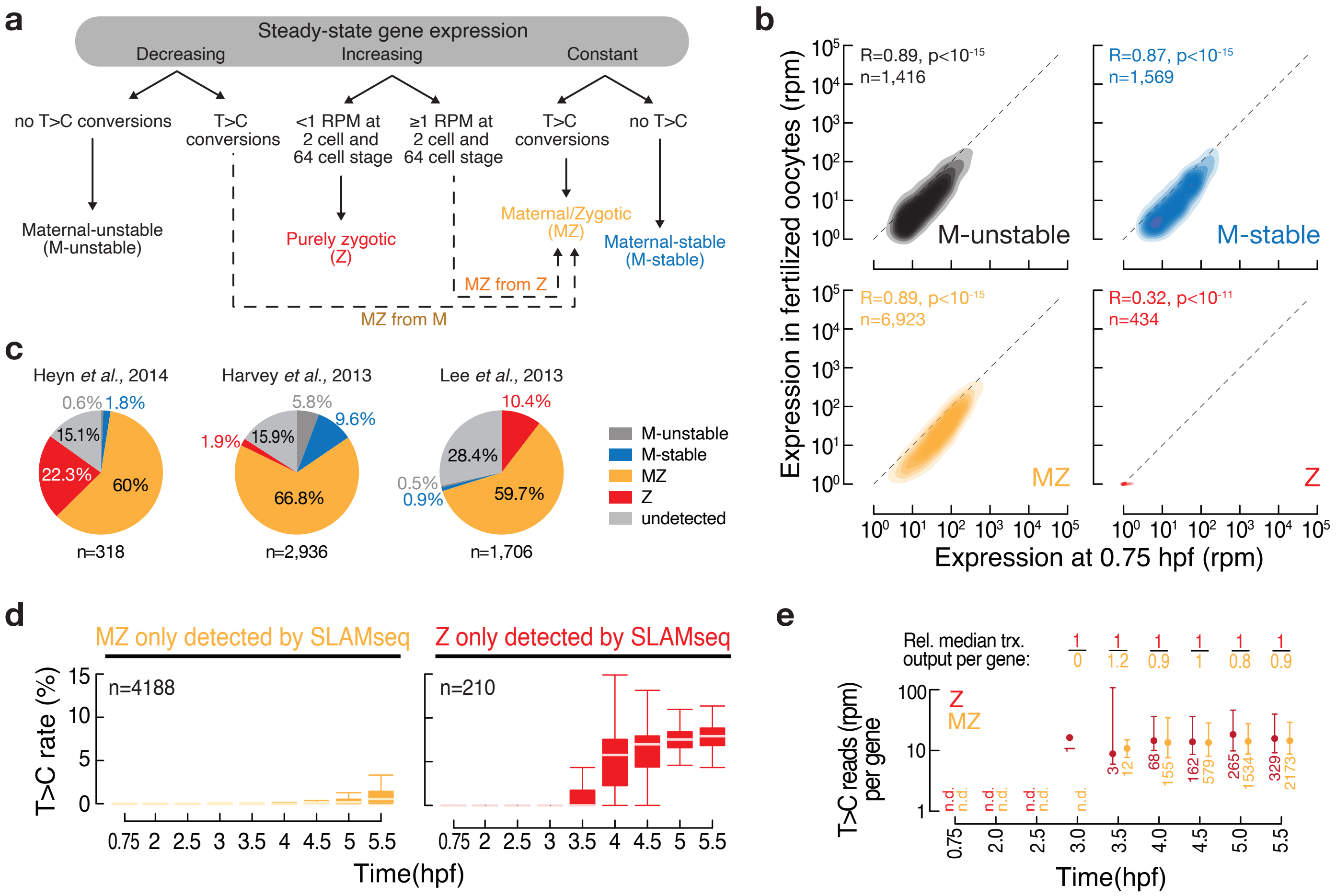

### Figure S5

Bhat et. al., Figure S5

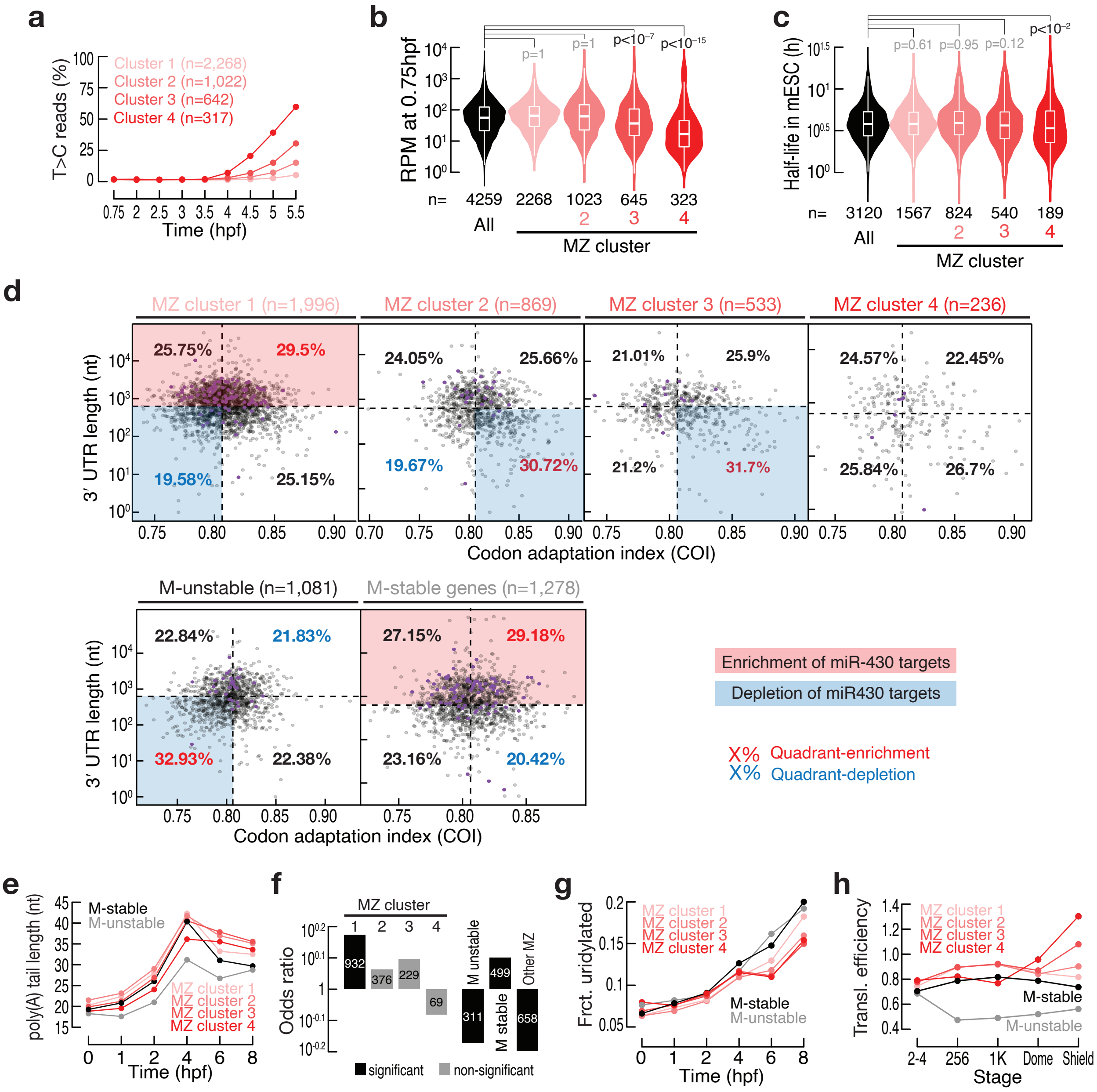
